# Biomimetic microstimulation of sensory cortices

**DOI:** 10.1101/2022.11.11.516221

**Authors:** Christopher Hughes, Takashi Kozai

## Abstract

**Background:** Intracortical microstimulation (ICMS) is an emerging approach to restore sensation to people with neurological injury or disease. Biomimetic microstimulation, or stimulus trains that mimic neural activity in the brain through encoding of onset and offset transients, could improve the utility of ICMS for BCI applications, but how biomimetic microstimulation affects neural activation is not understood. Stimulus induced depression of neural activity (decreases in evoked intensity over time) is also a potential barrier to clinical implementation of sensory feedback, and biomimetic microstimulation may reduce this effect.

**Objective:** We evaluated how biomimetic trains change the calcium response, spatial distribution, and depression of neurons in the somatosensory and visual cortices.

**Methods:** Calcium responses of neurons were measured in Layer 2/3 of visual and somatosensory cortices of anesthetized GCaMP6s mice in response to ICMS trains with fixed amplitude and frequency (Fixed) and three biomimetic ICMS trains that increased the stimulation intensity during the onset and offset of stimulation by modulating the amplitude (BioAmp), frequency (BioFreq), or amplitude and frequency (BioBoth). ICMS was provided for either 1-s with 4-s breaks (Short) or for 30-s with 15-s breaks (Long).

**Results:** BioAmp and BioBoth trains evoked distinct onset and offset transients in recruited neural populations, while BioFreq trains evoked population activity similar to Fixed trains. Individual neurons had heterogeneous responses primarily based on how quickly they depressed to ICMS, where neurons farther from the electrode depressed faster and a small subpopulation (1-5%) were modulated by BioFreq trains. Neurons that depressed to Short trains were also more likely to depress to Long trains, but Long trains induced more depression overall due to the increased stimulation length. Increasing the amplitude during the hold phase resulted in an increase in recruitment and intensity which resulted in more depression and reduced offset responses. Biomimetic amplitude modulation reduced stimulation induced depression by 14.6±0.3% for Short and 36.1±0.6% for Long trains. Ideal observers were 0.031±0.009 s faster for onset detection and 1.33±0.21 s faster for offset detection with biomimetic amplitude encoding.

**Conclusions:** Biomimetic amplitude modulation evokes distinct onset and offset transients, reduces depression of neural calcium activity, and decreases total charge injection for sensory feedback in brain-computer interfaces by lowering recruitment of neurons during long maintained periods of ICMS. In contrast, biomimetic frequency modulation evokes distinct onset and offset transients in a small subpopulation of neurons but also reduces depression in recruited neurons by reducing the rate of activation.

## Introduction

Electrical stimulation of the nervous system is one of the oldest and most widely used methods for evoking neural activity [1], [2]. This method has allowed for identification of biophysical properties of neurons [3]–[5] and the organization of the brain [6], as well as the restoration of function in people with neurological injuries or disease. Intracortical microstimulation (ICMS) has restored tactile sensation to the hands or arms of people with spinal cord injury [7]–[10] and restored visual percepts in people with blindness [11]–[18]. Tactile percepts range from natural sensations of “pressure” and “touch,” to less natural sensations of “tingle” and “buzz,” [7], [8] while visual percepts are dots or lines of light [14], [18]. Humans can also discriminate ICMS frequencies, however, limitations of ICMS include unnatural and electrode dependent percepts [10], [19].

In addition to lacking naturalness, percepts adapt, or decrease in intensity, under continuous ICMS and can completely extinguish within a minute [20]. Continuous somatosensation is critical for dexterity [21]–[23] and a continuous visual stream is necessary for naturalistic vision [24], [25], so that extinction of percepts may provide a critical barrier to sensory restoration. Sensory adaptation occurs for normal sensory input, but perceived intensity decreases gradually over time without becoming imperceptible and recovers on the order of seconds to minutes [26]–[31]. In contrast, ICMS induced depression of neuronal excitability (SIDNE) takes up to days to fully recover [32], [33]. SIDNE refers to changes in excitability *following* ICMS, but depression of neuronal calcium activity also occurs *during* ICMS in animals, which is stronger at higher frequencies [34]–[38]. We will distinguish between these three phenomena here: sensory adaptation (occurring in response to normal sensory input), SIDNE, and depression during continuous electrical stimulation (DCS). DCS is difficult to measure with traditional electrophysiology because of electrical artifact produced by ICMS but can be observed with high-resolution imaging of neural calcium activity. While mechanisms underlying sensory adaptation are not fully understood, various mechanisms have been proposed from single neuron dynamics involving changes in ion distribution to pooled population responses across many neurons [39]. SIDNE has been suggested to occur through accumulation of potassium ions in the extracellular space causing depolarization block [40] or by the production of toxic factors related to electrochemical potentials at the electrode interface [32], while DCS has been suggested to be a result of recruited inhibition or metabolic stress [34]. Exactly how sensory adaptation compares to SIDNE or DCS is unclear, but there seem to be clear differences in the magnitude and time course of these effects, so that ICMS-induced depression may provide a clinical barrier for continuous sensory restoration. Therefore, we were interested in knowing how dynamic parameter modulation of pulse trains might impact ICMS-induced depression of neurons.

As opposed to dynamic changes in ICMS, which have not been well-studied, higher amplitudes and frequencies of fixed ICMS are known to result in more DCS and adaptation [34]–[38]. Dynamic changes in amplitude and frequency may reduce depression by saliently encoding onset and offset and reducing the intensity of recruitment for extended periods of ICMS. For somatosensory ICMS, stimulus intensity has been linearly scaled to force detected on prosthetic hands by increasing ICMS amplitude, which improves robotic arm control [41] while visual ICMS has previously only provided short stimulus trains to represent simple features such as location, luminance, and shape [18], [42]. However, the brain does not linearly scale input: in sensory cortices, neurons respond strongly to the onset and offset of sensory input with modest firing in between [43]–[52]. Peripheral nerves and somatosensory cortices have strong onset/offset responses [43], [46], but so do retinal ganglion cells, optic nerves, and visual cortices, where offset of visual input results in up to 200 ms of heightened activity as measured by electrophysiology [45], [47]–[52]. Biomimetic ICMS that mimics cortical activity, by encoding strong onset/offset transients with low intensities in between, may improve evoked percepts and utility of continuous ICMS for BCI applications [53]–[56]. Biomimetic algorithms implement these dynamic changes in ICMS intensity [54], [57], which improve the naturalness and utility of peripheral nerve stimulation [58], [59]. Additionally, ICMS bio-inspired algorithms in the somatosensory cortex worked as well as linear algorithms with less charge injection for object identification [60], reduced artifact in the motor cortex [61], and improved amplitude discrimination [62] which could be beneficial for bidirectional brain-computer interfaces [63]. Similar algorithms have reduced large onset transients for high-frequency stimulation in the hippocampus and nerve [64], [65] and have reproduced natural activation patterns in the thalamus [66]. However, how biomimetic amplitude and/or frequency modulation might impact spatiotemporal neural activity in the cortex remains unknown.

Although spatiotemporal activation by biomimetic modulation in cortex has not been studied, biomimetic ICMS trains should be composed of parameter ranges relevant to clinical and rodent ICMS studies so that they are relevant to previous literature and clinical research. ICMS for somatosensation in humans and non-human primates (NHPs) has been limited to amplitudes of 0-100 µA, frequencies of 0-300 Hz, pulse durations of less than 1 ms, and train durations of less than 15 s [7], [8], [10], [67]–[69] which was based on previous animal work establishing safe and effective ranges for ICMS [70]–[73]. For vision, ICMS has consisted of short stimulus trains at high frequencies (50 pulses at 300 Hz) [18], [42]. In clinical bidirectional brain-computer interfaces, ICMS frequencies below 30 Hz are not used because individual pulses can be perceived as tapping, which does not accurately represent grasping or object contact [10]. Despite the use of high-frequency stimulation in clinical research [8], [10], [41], most excitatory neurons fire at low frequencies (average ∼8 Hz) [74]–[76]. Nevertheless, ICMS frequencies of up to 300 Hz are discriminable in humans and NHPs. Specifically, NHPs and humans can discriminate 100 Hz trains from 150 Hz trains and 100 Hz from 250 Hz trains with above 75% accuracy, respectively [69], [19]. Perceptually, higher frequencies tend to evoke more consistent “pressure,” “tingle,” and “touch” sensations [10]. In rodents, spatiotemporal activation changes with ICMS frequency [34], [35], [37]. Low frequency (10 Hz) ICMS evokes sustained activation of excitatory neurons, suggesting entrainment [35], [37]. However, high frequency (100-200 Hz) ICMS recruits the same number of neurons as 10 Hz, but as frequency increases, activation is followed by faster and greater depression of calcium responses. As a result, higher frequencies of ICMS (such as 130 vs. 250 Hz) evoke distinguishable rates and magnitudes of depression [34], [35], [37]. These studies demonstrate that different high frequencies of ICMS (50-250 Hz) can evoke distinguishable responses in clinical and animal studies and motivate the study of high frequency ranges for future applications of biomimetic encoding.

While different amplitudes and frequencies of ICMS can evoke distinct spatiotemporal profiles in rodents, how biomimetic modulation changes the spatiotemporal responses has not been previously studied. Studying the spatiotemporal activation of ICMS requires higher resolution readouts than 400 µm spaced Blackrock arrays typically used in humans and NHPs [7]–[9], [18], [42], [69]. Transgenic mouse models with high-resolution imaging have been useful for studying how ICMS parameters affect the spatiotemporal response of the brain and how ICMS parameters can change neural responses [34], [35], [37], [38], [77]–[79]. Biomimetic ICMS is being investigated in humans for somatosensory ICMS [60], [62], [80], but how biomimetic trains affect brain activation or depression is not understood. Measuring the neural response to ICMS in humans is difficult due to safety and ethical concerns over human transgenic manipulation and limitations of hardware and high-resolution imaging, challenges with deconvolving electrophysiological single-units from stimulation artifacts, and low throughput data given the limited number human subjects and their limited availability for testing. Therefore, we used transgenic GCaMP6s mouse models as a high throughput method to study mechanisms of biomimetic ICMS in Layer 2/3 of visual and somatosensory cortices.

Here, we expand previous work using high-resolution imaging in transgenic mouse models to investigate if biomimetic encoding of onset/offset through dynamic changes in amplitude and/or frequency would result in significant changes in the spatiotemporal activation and reduce depression compared to fixed-trains. Biomimetic amplitude modulation consistently increased the activity at the onset/offset of ICMS across 100% of recruited neurons. Biomimetic frequency modulation resulted in only a small subpopulation (1-5%) having onset/offset responses. Additionally, biomimetic amplitude and frequency modulation maintained greater excitability of neurons over long periods of continuous stimulation by reducing the intensity of recruitment during the hold phase. This work provides evidence that dynamic changes in ICMS amplitude and frequency can help reduce charge injection while enhancing onset/offset activity and maintaining excitability to repeated stimulation. Dynamic changes in frequency resulted in more subtle differences in activity than dynamic amplitude modulation within the frequencies evaluated, although reductions in frequency still reduced depression to continuous ICMS and reduced the overall charge injected. The findings of this study ultimately support the claim that dynamic modulation of stimulus amplitude and frequency can be an important tool in the modulation of neuronal activation for ICMS train and highlights the importance of considering nonlinear dynamics of ICMS parameters when designing electrical stimulation protocols for better control of neuronal activation.

## Methods

### Surgery and electrode implantation

Surgeries were performed on transgenic Thy1-GCaMP6s male (n=6) or female (n=6) mice (Jackson Laboratories, Bar Harbor, ME) >4 weeks of age weighing 25-35 g. Anesthesia was induced with ketamine and xylazine as described previously [34], [38]. Head-fixed mice were given rectangular craniotomies (2×3mm) over each visual cortex (n=8, long side 1 mm from sagittal and short side <1 mm from lambdoid suture) or somatosensory cortex (n=4, long side 1 mm from coronal and short side 1mm from sagittal suture), with a dental drill. We performed experiments in both visual and somatosensory cortex because we were interested in how the overall response might deviate between these two cortices given parallel work with ICMS for both vision and somatosensation restoration [7]–[10], [18]. Electrodes were generally targeted to the right hemisphere unless there was bleeding or injury. A bone screw was implanted in the left motor cortex for ground and reference. Single shank 3 mm long electrode arrays with 16 Iridium channels of 703 µm^2^ electrode sites spaced 50 µm apart (NeuroNexus Technologies, Ann Arbor, MI) were implanted at a 30° angle from the horizontal plane (Fig. 1A) using a Microdrive (MO-81, Narishige, Japan) to depths of 1000 µm. Electrodes spanned the first 800 µm of the electrode (Fig. 1A), so that the 1000 µm depth result in electrode sites spanning from ∼100 µm below the surface to ∼475 µm below the surface. Cross-correlation (xcorr) was used to determine similarities across cortices, sex, and trial (Supp. Fig. 1-3). Evoked activity with highly similar time courses returns high correlation values (>0.99). Electrode sites were activated prior to surgery based on published protocols [81]. Data within individual animals were all collected in a single session (of 3-5 hours).

**Figure 1:**
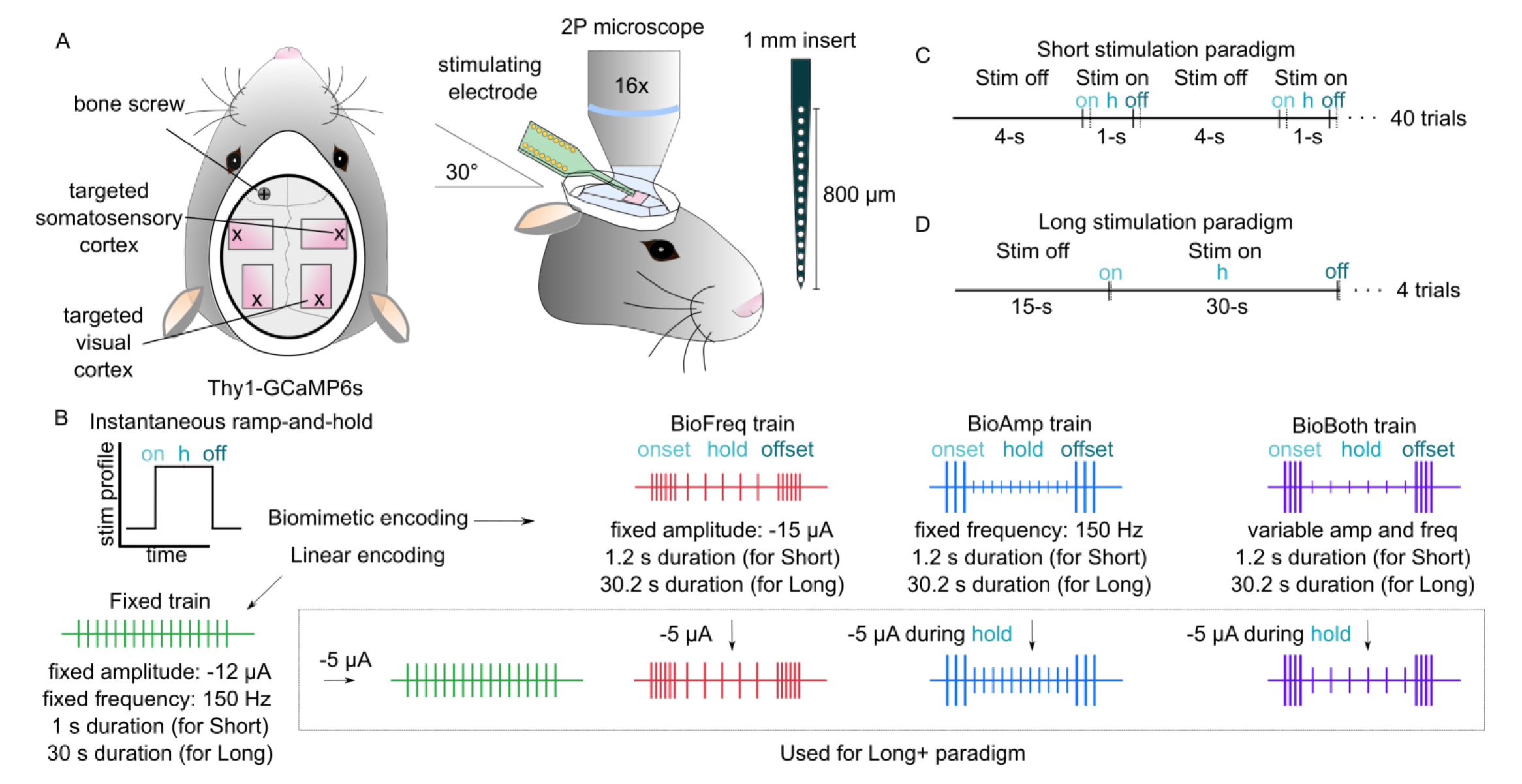
GCaMP6s mice received biomimetic microstimulation through Michigan probes with simultaneous calcium imaging. A) Electrodes were implanted at a 30° angle into the visual or somatosensory cortices of Thy1-GCaMP6s mice under anesthesia. A bone screw was placed into the skull for ground/reference. Two-photon imaging was used to image the open cranial window where ICMS was applied. Probes were inserted 1 mm so that all electrodes were in the cortex. B) Different ICMS trains were designed based on a transform from an instantaneous ramp-and hold function to either linearly encode the sensory input (Fixed) or to emulate the response of sensory cortex that either varied the amplitude (BioAmp), frequency (BioFreq), or both (BioBoth) for a pulse train with onset, hold, and offset phases. All 1-s trains were designed to be charge matched. For the Long+ trains we increased the amplitude by −5 µA during the hold phase for BioAmp and BioBoth and during the whole train for BioFreq and Fixed (shown in the box) C) For Short trains, we delivered 1-s (or 1.2-s) pulse trains followed by 4-s (or 3.8-s) breaks for 40 trials per block. Onset, hold, and offset phases are marked with “on,” “h”, and “off” and indicated by the vertical lines D) For Long trains, we delivered 30-s (or 30.2-s) pulse trains with 15-s (or 14.8-s) breaks for 4 trials per block.

### Electrical stimulation

A TDT IZ2 stimulator controlled by an RZ5D system (Tucker-Davis Technologies, Alachua, FL) applied current-controlled cathodic leading biphasic asymmetric pulses (200-μs cathodic, 100-μs interphase, 400-μs anodic), which matched human experiments [7]–[10]. Amplitudes in the range of 0-20 µA were used based on safety limits to reduce electrode/tissue damage [72], [73] and to be similar to ranges used in previous experiments in mice using Michigan probes [34], [35], [37], [77]. Frequencies of 0-200 Hz were used similar to ranges used in mice, NHPs, and humans [7], [8], [10], [20], [34]–[37], [67]–[69]. Only one electrode located in the imaging plane (120-200 µm below the surface) with an impedance <700 kOhms was used in each animal.

Stimulus trains were Short (1 s on, 4 s off), Long (30 s on, 15 s off), or Long+ (Long with −5 µA during the hold phase, Table 1). Short and Long trains were both used because Short trains match stimulation lengths used more commonly in studies in humans and NHPs for perception and psychophysics [7], [8], [10], [67]–[69]. For Short trains, the amplitude and frequency were balanced so that all trains injected the same amount of charge. In contrast, Long trains extended the hold phase to 30 s with the same 200 ms transients as the Short trains, resulting in differences in total charge injection between trains. Long trains better represent the network in an equilibrium state 15-20 s following initial recruitment [65], which can provide insight into the progression of excitation, inhibition, and disinhibition [34], [38], [65] and is one of the main reasons that Long trains are more commonly used in rodent studies [34], [35], [37], [38]. Furthermore, due to latencies in GCaMP activity [82], [83], longer stimulus profiles can provide more information about depression and dynamic intensity changes and provide more realistic lengths of ICMS for continuous sensory feedback applications. We additionally included Long+ trains (Long trains with larger hold amplitudes) to 1) determine to what extent low amplitudes during the hold phase drove the observed spatiotemporal effects of biomimetic modulation, 2) to determine if an ideal observer could discriminate two trains based on intensity differences (encoding intensity differences is important for stimulus discrimination), and c) to measure the effect of amplitude on the intensity and SIDNE/DCS over long periods of stimulation.

### Biomimetic stimulation parameters

Biomimetic profiles were designed to have high intensity “onset” and “offset” transients, as previously described for sensory input [54], [57]. All trains were designed to encode an instantaneous ramp-and-hold profile of either 1 s (Short) or 30 s (Long) (Fig. 1B). Ramp-and-hold profiles are common representations of force or intensity of stimulus input with an onset ramp, a hold period, and an offset ramp [43]. The rate of change of the ramps represents the speed at which the stimulus input goes from zero to the maximum intensity (where a steeper ramp represents a faster rate of change) and the hold period represents maintained contact or input at the maximal intensity. All pulse trains used here were designed to encode an instantaneous ramp-and-hold profile, where the hold period is achieved in 0 s (so the ramp is a vertical line) (Fig. 1B). An instantaneous ramp-and-hold could represent a mechanical indentation with a fast indentation rate or a visual input with fast increase to maximal luminance (i.e. a flash of light).

**Table 1:** Outline of the parameters used in each ICMS train. The 200-ms offset phase for each biomimetic train signaled the end of stimulus input which resulted in an additional 200 ms of ICMS compared to the Fixed train. All Short trains were symmetric and had equal total charge injection (360 nC, Table 1). For Long trains, consistent transient “onset” and “offset” periods (200-ms) were applied but with an extended “hold” phase (29.8-s), which resulted in unequal charge injection across trains, with lowest charge injection for BioBoth (1760 nC) and highest charge injection for Fixed trains (10800 nC). The Fixed trains did not have distinct onset and offset phases, so parameters are only specified for the hold phase

|  |  | Onset |  |  | Hold |  |  | Offset |  |  | Total charge (cathodic phase for all pulses, nC) |
| --- | --- | --- | --- | --- | --- | --- | --- | --- | --- | --- | --- |
|  |  | Amp (μA) | Freq (Hz) | Dur (s) | Amp (μA) | Freq (Hz) | Dur (s) | Amp (μA) | Freq (Hz) | Dur (s) |  |
| BioAmp | Short | -20 | 150 | 0.2 | -5 | 150 | 0.8 | -20 | 150 | 0.2 | 360 |
|  | Long | -20 | 150 | 0.2 | -5 | 150 | 29.8 | -20 | 150 | 0.2 | 4560 |
|  | Long+ | -20 | 150 | 0.2 | -10 | 150 | 29.8 | -20 | 150 | 0.2 | 8880 |
| BioFreq | Short | -15 | 200 | 0.2 | -15 | 50 | 0.8 | -15 | 200 | 0.2 | 360 |
|  | Long | -15 | 200 | 0.2 | -15 | 50 | 29.8 | -15 | 200 | 0.2 | 4560 |
|  | Long+ | -20 | 200 | 0.2 | -20 | 50 | 29.8 | -20 | 200 | 0.2 | 6080 |
| BioBoth | Short | -20 | 200 | 0.2 | -5 | 50 | 0.8 | -20 | 200 | 0.2 | 360 |
|  | Long | -20 | 200 | 0.2 | -5 | 50 | 29.8 | -20 | 200 | 0.2 | 1760 |
|  | Long+ | -20 | 200 | 0.2 | -10 | 50 | 29.8 | -20 | 200 | 0.2 | 3200 |
| Fixed | Short | N/A | N/A | N/A | -12 | 150 | 1 | N/A | N/A | N/A | 360 |
|  | Long | N/A | N/A | N/A | -12 | 150 | 30 | N/A | N/A | N/A | 10800 |
|  | Long+ | N/A | N/A | N/A | -17 | 150 | 30 | N/A | N/A | N/A | 15300 |

Fixed-parameter trains were delivered at a constant amplitude and frequency (Fixed) across the entire train. Biomimetic trains provided high-intensity “onset” and “offset” transients (200 ms in length) with high amplitude modulation (BioAmp), frequency modulation (BioFreq), or both (BioBoth) with a 25% low-intensity “hold” phase in between (Fig. 1B). The offset phase for biomimetic encoding occurred for 200 ms following the end of the encoded stim profile (either 1 s or 30 s) so that the offset signal began when the instantaneous ramp-and-hold ended, signaling the end of the stimulus input. 200 ms was chosen and placed directly after the onset or offset because the time course of onset and offset activity recorded previously in both somatosensory and visual cortices is about 200 ms [43], [47], [48]. Informed decisions were made so that the tested trains could accurately evaluate the outlined questions regarding how biomimetic (dynamic) parameter modulation affected the spatiotemporal response and SIDNE/DCS of recruited neurons. We chose the Fixed train to be reasonably in the middle of the parameters used here (medium amplitude and frequency) and small variations in these parameters are unlikely to change the overall effect, which is validated by the small changes observed for Long vs. Long+ trains here. Train order was randomized to remove ordering effects. Data from 12 mice are presented (Supp. Table 1).

### Two-photon imaging

A two-photon laser scanning microscope (Bruker, Madison, WI) with a 16X 0.8 numerical aperture water immersion objective lens (Nikon Instruments, Melville, NY) and an OPO laser (Insight DS+; Spectra-Physics, Menlo Park, CA) tuned to a wavelength of 920 nm were used. Images were recorded at 30Hz and 1.52× optical zoom with 512×512 pixel resolution with a total coverage of 543.2×543.2 µm (Supplemental Movies 1-8).

### Image processing

Cells were manually outlined using 2D projections of the standard deviation and grouped averages of fluorescence using ImageJ (NIH). In animals with no significant shifts in the imaging plane across trials, a consistent region-of-interest (ROI) map was used to make comparisons of individual ROIs across trains. ROIs represent putative neurons, and are referred to as such throughout the text. Mean fluorescence over time of each ROI was extracted and imported to MATLAB (Mathworks, Natick, MA). All fluorescence data was filtered using a 15-sample Gaussian temporal filter to remove noise artifact. Fluorescent intensity was normalized to the baseline (dF/F0) at the beginning of the trial (3 s for Short, 14 s for Long). Neurons were parsed into active and inactive populations with a threshold of +3 standard deviations of baseline. For Short trains, onset was 0 to 1 s after ICMS start and offset was from 1 to 2 s. For Long trains, onset was 0 to 3 s after ICMS start, hold was from 3 to 30 s, and offset was from 30 to 33 s. Time to peak was calculated as the time to achieve the peak of activation following the start of the onset. Time to baseline was calculated as the time to achieve 0.5x the standard deviation of the baseline following the start of the offset.

### Classifying individual neurons responses

Neurons were divided into groups, or “classes,” based on temporal characteristics of the calcium evoked responses (Supp. Fig. 4). Neurons were first categorized into two categories; 1) “Modulated” if they showed substantial increases in the onset and offset transient (with two peaks), or 2) “Unmodulated” if no offset transient could be detected. For Short BioAmp and BioBoth trains, all neurons were identified as “Modulated.” If the first peak was greater than the second peak by at least 0.5x standard deviations of the noise, then the neuron was identified as “Onset modulated (Onset).” If the second peak was greater, the neuron was identified as “Offset modulated (Offset).” If the peaks were not different by more than 0.5x standard deviations of the noise, then the neuron was identified as “Dual modulated (Same).” Short BioFreq trains were identified to have a small number of neurons (<5%) that were “Modulated” by the onset and offset transients.

Neurons that did not have an Offset period greater than the Hold period were classified “Unmodulated.” All neurons for Fixed trains and most neurons for BioFreq trains were identified as Unmodulated. Unmodulated neurons were divided based on their rate of depression. For Short trains, neurons that depressed quickly (<85% of max at 1 s) were labeled “Rapidly Adapting” (RA), those that depressed more slowly (<95% of max at 1 s) were labeled “Slowly Adapting” (SA), and those that did not depress (>95% of max at 1 s) were labeled “Non-adapting” (NA).

Similarly, for Long and Long+ BioAmp and BioBoth trains, all neurons were identified as “Modulated” neurons and were subcategorized into “Onset Modulated (Onset),” “Same Modulated (Same),” and “Offset Modulated (Offset),” classes. Additionally, some neurons that had elevated activity during the hold period were separated as “Slowly Adapting” (SA) or “Non-Adapting” (NA) classifications. SA neurons had a maximum intensity in the second half of the hold period that was less than the first half by at least 0.5x standard deviations. Otherwise, neurons were classified as Non-Adapting (NA).

Long and Long+ BioFreq and Fixed trains also had SA and NA responses, but also had “Rapidly Adapting” (RA) responses. RA neurons had <50% of their max peak after only 10 s of stimulation, while SA neurons had >50% of their max peak at 10 s of stimulation. We used different definitions for RA and SA for Short and Long trains because of the different magnitudes of depression that occurred given the length of stimulation. However, we found that the classifications of individual neurons for Short and Long trains were strongly correlated, indicating that these phenomena represent a consistent response property of individual neurons despite the different definitions (Supp. Fig. 7).

### Adaptation indices

Adaptation indices (*AI*s) of fluorescent intensity for continuous ICMS were calculated with:

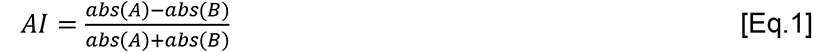

where *A* represents the mean activity in a neuron during the first half of the evoked response (0-1 s for Short, 0- 16 s for Long) and *B* represents the mean activity in the second half of the evoked response (1-2 s for Short, 16- 32 s for Long). For cross-trial adaptation, the mean was calculated across all active neurons within a trial and *A* is the first trial while *B* is the last trial. An *AI* of 1 would demonstrate complete adaptation, 0 no adaptation, and −1 complete facilitation (increase in activity).

### Ideal observer analysis

An ideal observer classifies stimuli determined by the arguments of maxima from Baye’s formula (i.e. Bayesian classification) [84]. Ideal observer analysis has been used previously to classify natural-and ICMS-evoked sensory responses with voltage-sensitive dyes [85], [86]. Here, Bayesian classifiers were trained on the average fluorescent activity across neurons in response to Long trains. Stimulus amplitude (which is known to increase neural recruitment and intensity [36], [77], [87], [88]) also results in better detection, faster reactions, and higher intensities of percepts [7], [10], [62], [68], [69], [89]. Therefore, by quantifying the speed and intensity of the calcium evoked response, the ideal observer analysis is reasonably quantifying the likelihood that the stimulus profiles would modulate the salience of the stimulus input and the detection time. However, because of latencies in GCaMP signal as well as the need for upstream/downstream cortical integration of signal for behavior, the times to detection provided here are not representative of real perceptual detection times. While this analysis then is informative about the potential utility of biomimetic amplitude modulation to improve ICMS utility, changes in detection will need to be studied with behavioral experiments in animals or humans.

Times of onset and offset were determined with biomimetic and fixed encoding models. These models and their application are described in Supp. Fig. 8A. Bayesian classifiers, using Bayes’ theorem to assign observations to the most probable class [90], were trained using the “fitcnb” function in MATLAB and classification scores were returned using the “resubPredict” function. A simpler thresholding method was also applied, in which changes in state were determined with a threshold of +3 SDs baseline (Supp. Fig. 8C,D).

For discrimination of stimulus intensity, activity during the hold phase was adjusted with Eq. 2 (i.e. the intensity over time was adjusted based on the function describing intensity-driven adaptation so that classification was based on the overall intensity and not impacted by adaptation) and each classifier was trained to distinguish low amplitude trains from paired high amplitude trains. Accuracy of the classifiers were combined into single columns for each train across animals and significant differences were calculated between accuracies for each train.

### Statistics

All statistical analyses were conducted in MATLAB. All data were compared for significant differences at an α of 0.05 using ANOVA followed by Tukey’s HSD to identify pairwise differences and account for multiple comparisons unless otherwise noted. Significance is indicated in figures with * for *p*<0.05, ** for *p*<0.01, and *** for *p*<0.001. Violin plots were generated by estimating the kernel density using ksdensity() and cutting off values at min/max (for example, a lower limit of 0 and upper limit of 543 µm for distance based on the imaging window) possible values [91]. All means are reported ±SE.

## Results

### Investigating the mechanisms of biomimetic stimulation parameters

Two-photon imaging in transgenic mice allows for measuring cellular resolution spatiotemporal responses to intracortical microstimulation (ICMS). These measurements allow for high throughput investigation of neural mechanisms underlying ICMS, which are currently difficult to evaluate in humans [34], [35], [37], [38], [77], [79]. Biomimetic ICMS aims to reproduce naturalistic activation of the brain with trains that mimic the cortical onset and offset response to sensory input through dynamic modulation of stimulus parameters [53], [54], [62]. Here, we encoded a 200 ms high amplitude and/or frequency segment into ICMS trains to drive a post-stimulus offset response to mimic activity following natural sensory cessation. The cortical response to biomimetic ICMS has not been studied previously, so how biomimetic trains activate cortical neurons remains a gap in knowledge. Here, we measured the response of excitatory neurons in Layer 2/3 of somatosensory and visual cortices of GCaMP6s mice with biomimetic and fixed-parameter ICMS trains. Specifically, we applied dynamic changes in amplitude and/or frequency in clinically relevant ranges and measured spatiotemporal activation of recruited neurons, stimulus-induced depression of neuronal excitability (SIDNE), and depression during continuous stimulation (DCS). We used both Short (1 s) and Long (30 s) trains to study how different lengths of ICMS affect patterns of activation, given the large amount of literature on perception with Short ICMS trains in humans and NHPs [7], [8], [10], [62], [68], [69] and on network equilibrium state when continuous ICMS is sustained with Long trains in rodents [34], [35], [37], [38], [65]. Individual neurons were categorized based primarily on modulation to offset transients and rates of ICMS induced depression. Biomimetic modulation significantly changed the responses of recruited populations and individual neurons as well as DCS and SIDNE.

### Biomimetic amplitude modulation evokes distinct onset and offset transients in recruited populations more than biomimetic frequency modulation

Biomimetic profiles are designed to mimic the onset and offset population response of cortical neurons to sensory input through amplitude or frequency modulation. Increasing ICMS amplitude increases the number of neurons activated, primarily by increasing the density of recruited neurons near the electrode [77], [87], [88]. Increasing ICMS frequency can increase activity over short durations, but generally causes depression of activity over extended durations [34]–[37]. We customized ICMS trains to modulate both amplitude and frequency to encode onset/offset of ICMS input. To evaluate how well each train evoked onset/offset responses, we looked at the average population response for each train type. We hypothesized that biomimetic modulation of pulse trains would substantially increase the onset/offset response of the population of neurons recruited by ICMS compared to fixed parameter stimulation.

#### Biomimetic amplitude modulation results in distinct onset and offset responses, increased intensity, and faster times to peak for Short trains

We first investigated Short ICMS trains (1 or 1.2 s on) which are typical of previous perceptual experiments in humans [7], [8], [10], [67]–[69]. Short ICMS trains evoked calcium activity for 3.04±0.009 s following onset (Fig. 2A,B). Biomimetic amplitude modulation (BioAmp and BioBoth trains) resulted in distinct onset/offset transients (Fig. 2C,D), but biomimetic frequency modulation alone (Fig. 2E) did not evoke distinct onset/offset transients and the response for BioFreq trains closely resembled the response to Fixed trains (Fig. 2F). BioAmp/BioBoth Short trains resulted in faster times-to-peak (BioAmp=0.40±0.001 s, BioBoth=0.40±0.002 s vs. BioFreq=0.75±0.009 s, Fixed=0.50±0.004 s, *p*<0.001, Fig. 2G). Faster times-to-peak could indicate decreased latency to stimulus detection, but may also be due to calcium latencies [82], [83]. BioAmp/BioBoth Short trains also resulted in slower recovery to baseline (BioAmp=2.31±0.024s, BioBoth=1.97±0.027s vs. BioFreq=1.91±0.032s and Fixed=1.77±0.043s, *p*<0.001, Fig. 2H) which could reduce the ability to deliver stimulus trains in quick succession. Average evoked activity across all Short trains recovered to baseline in less than 2.5 s after the end of ICMS (Fig. 2H), which indicates that these ICMS trains can be delivered within 4 s of one another without directly impacting the evoked calcium activity of the following train. Together, biomimetic amplitude modulation evoked strong onset/offset responses for Short trains compared to biomimetic frequency modulation. Biomimetic amplitude modulation also resulted in faster times to peak activation which could represent decreased time to stimulus representation, although biomimetic amplitude modulation also resulted in slower recovery to baseline which could decrease the temporal resolution over which trains can be delivered.

**Figure 2:**
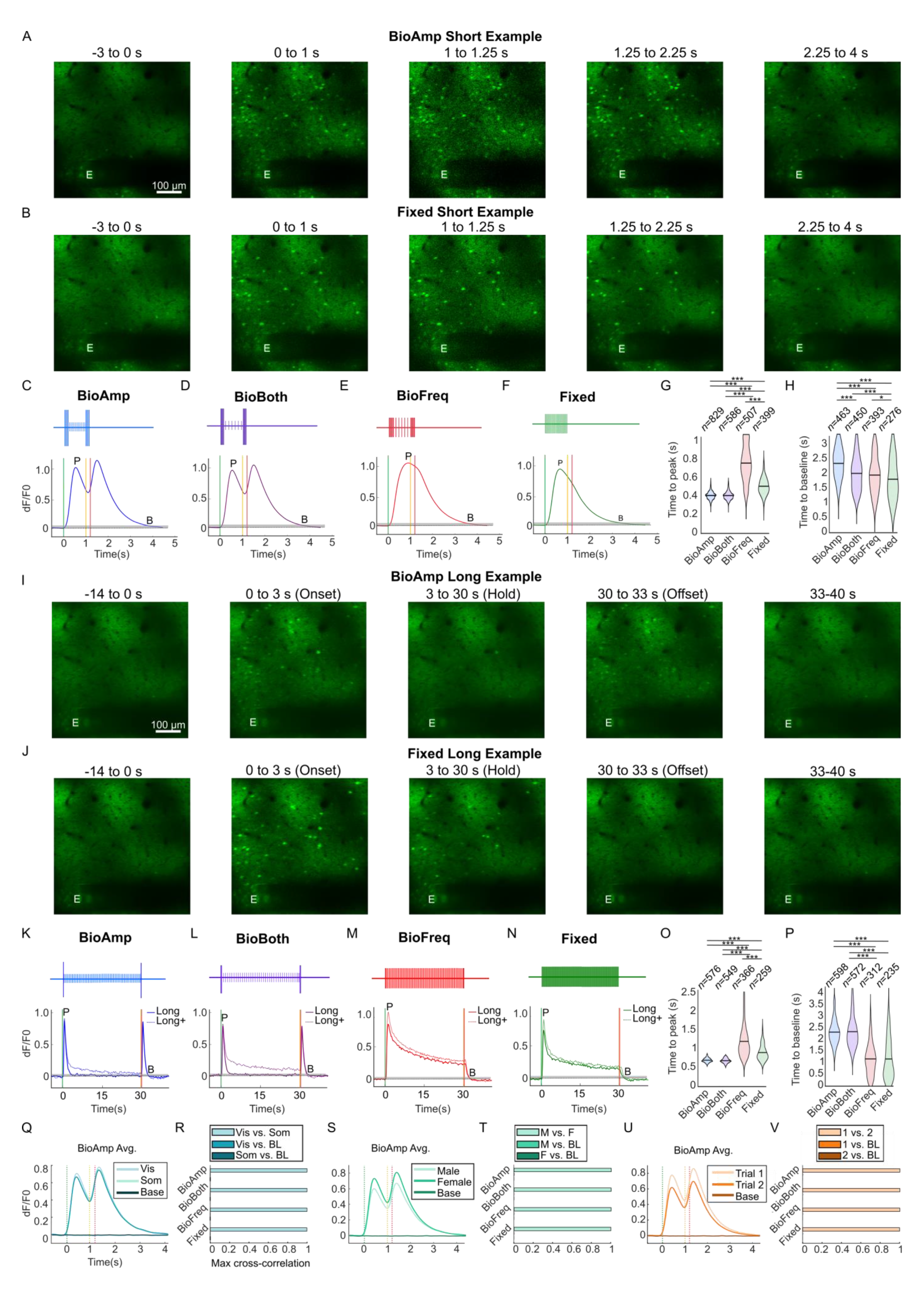
Biomimetic amplitude modulated trains evoke strong onset and offset transients and weak hold activation in recruited populations, while biomimetic frequency or fixed trains evoke strong activity that depresses over time and these responses are consistent across sensory cortices and animal sex. A-B) Examples of 2P images of evoked activity for a Short BioAmp (A) and Fixed (B) train. For illustration, images shown were smoothed using singular value decomposition, but were smoothed with a temporal Gaussian filter for analyses. The “E” marks the stimulated electrode location. Scale bar=100 µm wide, 10 µm thick. C-H) Short train responses (*n*=12 animals) C-D) BioAmp and BioBoth trains evoked increases in the intensity during the onset and offset phases. Each trace shows the average activity across all neurons. Vertical dotted lines mark the onset start (green), offset start (yellow), and offset end (red). The grey box represents 0.5x the standard deviation of the average baseline across all neurons. The letters “P” and “B” mark the location of the time-to-peak and time-to-baseline for the average trace. E-F) BioFreq and Fixed trains evoked similar profiles. G) BioAmp and BioBoth trains achieved a faster time to peak than the BioFreq and Fixed trains. The size of the violin represents the kernel density estimate of the sample, the black bar shows the mean. Outliers more than three scaled median absolute deviations from the median were excluded for illustration. H) Fixed trains returned to baseline faster than the other trains. I-J) Examples of 2P images for Long BioAmp (I) and Fixed (J) trains. K-P) Long (solid lines, *n*=9 animals) and Long+ (dotted lines, *n*=7 animals) trains. K-L) BioAmp and BioBoth trains reduced activation during the hold phase, maintaining excitability for the offset transient. M-N) BioFreq and Fixed trains maintained strong activation during the hold phase driving depression of firing activity. O-P) BioAmp and BioBoth trains achieved peak faster and baseline slower than Fixed and BioFreq trains. Q-T) Cross-correlation analysis showing the Short BioAmp trains as an example. The average response is essentially identical across cortices (Q), sex (S), and trials (T). Cross-correlation of the average response across cortices (R), sex (T), and trial (V) demonstrate essentially identical response profiles, with >0.99 correlation. Note that the correlation with baseline activity is near zero for all comparisons, making these bars barely visible in the bar plots (R,T,V).

#### Biomimetic amplitude modulation resulted in weak activation during the extended hold phase for Long trains, while frequency modulation resulted in elevated and depressing activity

In the 1 or 1.2 s Short ICMS trains, the evoked neuronal population activity does not have time to reach an equilibrium of excitation/inhibition or to demonstrate substantial depression during continuous stimulation (DCS), especially because of latencies in GCaMP [82], [83]. Therefore, to gain insight on competing excitatory and inhibitory activity and how different encoding models impact DCS, we extended each train to 30 seconds (Long) (Fig. 2I,J). We additionally tested a Long train with increased amplitudes (−5 µA; Long) during the hold period to better understand how amplitude affects the population activity profile, and particularly the offset response. These Long and Long+ trains evoked similar responses to Short trains but with extended hold phases (Fig. 2K-N). Long+ trains evoked significantly higher mean intensities than Long trains during the stimulation period (*p*<0.001) except for the Fixed train (*p*=0.21) primarily through increasing the intensity of activation during the hold period. While the Fixed Long+ train appeared to have initially higher intensity of activation, the strong depression induced by the higher amplitude made the overall evoked intensity not significantly different from the Fixed Long train (Fig. 2N). Long BioAmp/BioBoth trains weakly activated neurons during the hold phase (resulting in weak depression of neural calcium activity, Fig. 2K,L), while Long BioFreq and Fixed trains evoked strong and depressing responses in recruited neurons (Fig. 2M,N). BioAmp/BioBoth Long trains achieved much faster times to peak, similar to the Short trains (*p*<0.001, Fig. 2O). Evoked activity in Long trains recovered to baseline in less than 4 s (average 2.5 s) after the end of ICMS, similar to Short trains (Fig. 2P). BioFreq and Fixed trains returned to baseline more quickly, likely because the calcium activity had depressed substantially by the end of the ICMS train compared to the peak offset response of BioAmp/BioBoth trains (*p*<0.001, Fig. 2P). Together, Long and Long+ biomimetic amplitude modulation but not frequency modulation substantially altered the population response relative to Fixed trains. Long and Long+ trains evoked similar activation patterns to Short trains, but with much stronger DCS that occurred across the 30-s stimulation period.

#### ICMS profiles evoked similar population responses across cortices, animal sex, and trials

Finally, we were interested in knowing if other variables in our experiments contributed to the ICMS evoked responses to inform our own interpretation as well as future experiments and clinical applications. We found using cross-correlation analysis that there were no detectable differences in timing and shape of the population response evoked across visual vs. somatosensory cortices, males vs. females, and repeated trials (corr. Coeff > 0.99, Fig. 2Q-T, Supp Fig. 1-3). For the rest of this paper, we combine across cortices and animal sex to evaluate the effects of each train type on evoked responses.

Overall, we found that BioAmp and BioBoth trains encoded strong onset/offset responses while BioFreq and Fixed trains evoked continuous activity that depressed over time for all train lengths, which was consistent across cortices and animal sex. BioAmp and BioBoth trains achieved faster times to peak which could potentially indicate decreased time to cortical stimulus representation, although decreased times to peak may also be due to calcium latencies. Additionally, calcium activity returned to baseline in less than 4 s for all trains, indicating that trains can be delivered within 4 s of each other without calcium activity from one train overlapping with the calcium response of the next train. These findings are important because they clarified that biomimetic amplitude modulation could strongly modulate recruited neural populations to the onset/offset, similar to naturalistic activation patterns of the brain, and can be used similarly across different sensory cortices for sensory restoration.

### Biomimetic amplitude modulation increases total neurons recruited and the strength of individual responses during offset with small changes in distance from the electrode

Having found that biomimetic amplitude modulation evoked strong onset/offset transients in the population response, we wanted to know if the mechanisms of onset/offset activation were driven by larger recruitment of neurons or greater intensities of activation in recruited neurons. Additionally, we wanted to know if a greater total number of neurons were recruited near the electrode or new neurons were recruited farther from the electrode. Previous computational work demonstrated that higher ICMS amplitudes can increase the density of activation by recruiting more neural processes, but average soma distances do not change substantially [87], which agrees with results from experimental work [34], [35], [37], [38], [77]. We hypothesized that the high amplitude onset/offset transients of BioAmp/BioBoth trains would increase total recruitment and intensity of neurons without substantial changes in the distance of recruited soma from the electrode. The BioFreq/Fixed trains would have greater recruitment during the hold phase that would depress over time. Long trains verified the average onset/offset phases lasted 3 s (and the return to baseline took on average 2.5 s), which was longer than the 800 ms hold period in the Short trains. Thus, Short trains could not be meaningfully divided into phases and neurons recruited, mean intensity, and mean distance were all calculated from the whole activation profile (0-2 s). Long and Long+ trains were able to be divided into phases (onset, hold, and offset) because of the longer durations.

#### Total neuron recruitment with biomimetic amplitude modulation is substantially increased during offset and substantially reduced during the hold phase

To better understand how dynamic changes in parameters resulted in changes in population activity, we assessed how different train types, durations, and amplitudes affected the total recruitment of neurons. Short trains were first tested based on their relevance to human perceptual work. Although differences were not significant between trains (*p*=0.15), BioAmp trains evoked activity in the most neurons (active neurons=58±9%) and Fixed trains evoked activity in the fewest neurons (31±8%), representing a ∼27% difference (Fig. 3A). This result implied that recruitment was similar across train types, but the strong onset/offset responses contained within the BioAmp/BioBoth trains may dominate the recruitment of neurons over short periods.

**Figure 3:**
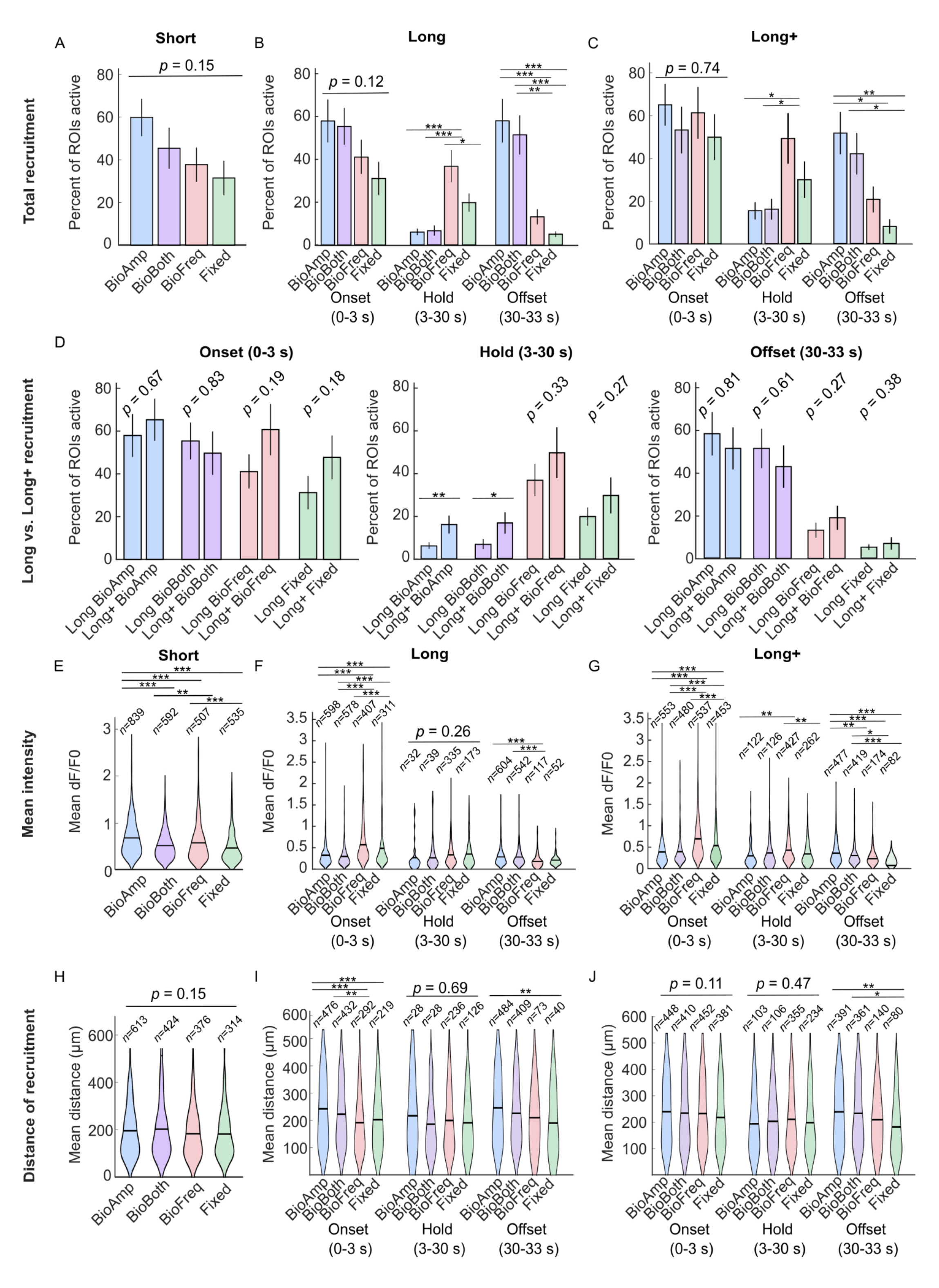
Biomimetic amplitude modulated trains evoke activity in more neurons at higher intensities during the offset phase with small but significant increases in distance, while Fixed and BioFreq trains evoke activity in more neurons at higher intensities during the hold phase with no significant differences in distance. A-C) Bar plots showing total recruitment of neurons for different train types, durations, and amplitudes of ICMS. Biomimetic amplitude modulation increases the percent of active neurons for A) Short trains (*n*=12 animals) and increases the percent active in the offset phase and decreases the amount of neurons active during the hold phase for B) Long (*n*=9 animals), and C) Long+ trains (*n*=7 animals). The bars for % active show the mean while the error bars show the standard error. D) Bar plots showing how recruitment of neurons was different between Long and Long+ trains. Higher amplitudes increased recruitment during the onset and hold periods for all trains. This decreased the excitability of neurons, resulting in a decrease in the offset response for BioAmp and BioBoth trains. E-G) Violin plots showing the mean evoked intensity of recruited neurons for different train types, durations, and amplitudes of ICMS. Biomimetic amplitude modulation increases the mean intensity for E) Short trains and increases the mean intensity during the offset phase, but decreases mean intensity during the onset and hold phase for F) Long and G) Long+ trains. The size of the violin represents the kernel density estimate of the sample, the black bar shows the mean. H-J) Violin plots showing the mean distance from the electrode of recruited neurons for different train types, durations, and amplitude of ICMS. Biomimetic amplitude modulation results in H) no significant changes distance for Short trains and I) increases in the average distance of recruitment during and offset for Long trains and J) increases in the average distance for the onset and offset for Long+ trains.

However, results from the Short trains did not reveal how recruitment changes within each phase (onset, hold, offset) due to the overlap of the 2.5 s calcium activity decay. To investigate how recruitment changes within the onset, hold, and offset phases, responses to Long trains were assessed. Although differences were not significant between trains during the onset phase (*p*=0.12), BioAmp and BioBoth trains evoked activity in more neurons (58±10% and 55±9%) than BioFreq and Fixed trains (41±8% and 31±8%), representing a 14-27% difference in recruitment (Fig. 3B left). This non-significant result might suggest that the transition from rest to ICMS onset drives an equally strong onset response regardless of train type. However, differences were significant when combining across BioAmp/BioBoth and BioFreq/Fixed (*p*=0.022) indicating that the lack of significance may have been potentially due to a weak effect relative to the sample size (*n*=9), so that BioAmp/BioBoth trains may evoke stronger onset activation. Higher amplitude transients then may increase recruitment of neurons, as expected [87], [88], [92]. During the hold phase, BioAmp and BioBoth Long trains recruited much less neurons overall (active neurons=2.9±0.9% and 3.9±1.5%) than BioFreq and Fixed Long trains (active neurons=34±7.3% and 17±3.9%, *p*<0.001, Fig. 3B center). The 75% reduction in amplitude then reduced the number of neurons recruited, as expected. We next looked at the offset phase to understand if the offset transient results in increased recruitment following long bouts of ICMS. During the offset phase, BioAmp and BioBoth Long trains evoked activity in more neurons (active neurons=58±10% and 51±9%) than BioFreq and Fixed Long trains (active neurons=12±3.5% and 4.6±1.2%, *p*<0.001, Fig. 3B right). Together, BioAmp/BioBoth trains were able to maintain excitability of neurons for the offset (achieving a similar peak to the onset phase) by decreasing recruitment during the hold phase, while the Fixed/BioFreq trains continuously evoked strong recruitment which resulted in gradual depression of neurons and decreased recruitment for the offset.

The reduced amplitude during the hold phase resulted in less recruitment of neurons, which may have allowed for DCS recovery and enabled the encoding of an offset response. However, to clarify if amplitude was solely responsible for changes in recruitment, we examined the sensitivity of the offset response to the hold amplitudes using Long+ trains. Long+ trains overall had similar trends to Long trains, with Fixed and BioFreq recruiting more neurons during the hold and BioAmp and BioBoth recruiting more neurons during the offset (Fig. 3C). We next compared the results of Long and Long+ trains directly (Fig. 3D). In most phases, recruitment was not significantly different between Long and Long+ trains, although Long+ trains generally evoked greater recruitment during the onset and hold phases (Fig. 3D left, center). Only the Long+ BioAmp and BioBoth trains evoked significantly more recruitment during the hold phase. Although differences were not significant, BioAmp and BioBoth Long+ trains recruited less neurons during the offset than Long trains (Fig. 3D right). The increased recruitment during the onset and hold phase may have resulted in less excitability for the offset phase. Together, amplitude modulation had a strong effect on the recruitment and corresponding depression of the recruited neural populations. Long+ trains may have reduced the offset response by maintaining greater recruitment during the hold phase.

#### Mean intensity of activation of recruited neurons with biomimetic amplitude modulation is also substantially increased during offset and substantially reduced during the hold phase

Given that amplitude modulation reduced the number of recruited neurons during the low amplitude hold phase, we wanted to examine if dynamic modulation also resulted in lower intensities of activation in individual neurons. Therefore, we next investigated how the mean activity changed across recruited neurons in clinically relevant ranges of amplitude and frequency modulation. Short BioAmp trains evoked the highest intensity of activation (mean dF/F0=0.69±0.01, *p*<0.001, Fig. 3E) and Short Fixed trains evoked the lowest activity (mean dF/F0=0.46±0.009), representing a ∼33% difference in mean intensity. This large difference in intensities suggests that the high amplitude onset/offset transients recruit stronger intensities in individual neurons over short durations of ICMS.

To clarify if increased intensity was due to more intense activation during onset/offset, we next investigated how intensity changed across phases with Long trains. BioFreq and Fixed Long trains elicited a significantly stronger onset mean intensity (dF/F0=0.60±0.022 and 0.49±0.028; *p*<0.05) compared to BioAmp and BioBoth Long trains (dF/F0=0.31±0.01 and 0.28±0.01). The higher dF/F0 activity of BioFreq and Fixed Long trains was due to greater sustained activity during this Onset defined period (0-3 s) whereas BioAmp/BioBoth trains rapidly decreased towards baseline after cessation of the onset transient (Fig. 3F left). Interestingly, during the hold phase, the calcium activity of active neurons was not significantly different between BioAmp/BioBoth (mean dF/F0=0.27±0.06 and 0.27±0.05) and BioFreq/Fixed trains (mean dF/F0=0.33±0.02 and 0.36±0.02, *p*=0.26, Fig. 3F center). BioAmp/BioBoth trains then recruit significantly fewer neurons during the hold period than BioFreq/Fixed trains (*p*<0.05; Fig. 3B), but neurons that remained active during the hold period had similar intensities across trains, likely because DCS driven by Fixed and BioFreq trains resulted in lower intensities. During the offset phase, BioAmp/BioBoth trains evoked significantly higher intensities (mean dF/F0=0.35±0.01 and 0.35±0.01) than BioFreq trains (dF/F0=0.19±0.01, *p*<0.001, Fig. 3F right) and non-significant increases over Fixed trains (0.23±0.02), representing a 34-46% difference in intensities, similar to changes in recruitment reported in the previous section. BioAmp/BioBoth trains then can modulate both the recruitment and intensity of activation of individual neurons, so that lowering the amplitude during the hold phase reduces intensity of activation of neurons likely allowing for reduced depression and strong offset responses.

To understand if higher recruitment during the hold period changed the mean responses during offset, we next looked at Long+ trains. Long+ trains similar to Long trains had higher evoked intensities during onset for BioFreq/Fixed trains and higher evoked intensities for BioAmp/BioBoth trains during offset (*p*<0.001, Fig. 3G left). However, the BioFreq train had significantly higher intensity than the BioAmp and Fixed trains during the hold period, suggesting that the lower frequency content of the BioFreq train at higher amplitudes resulted in less depression (*p*<0.01, Fig. 3G center). Additionally, more significant differences existed between the offset responses across Long+ trains than across Long trains, suggesting that the higher amplitude increased differences in depression between trains (*p*<0.001, Fig. 3G right). Together, biomimetic amplitude modulation resulted in changes to the total recruitment of neurons as well as the intensity evoked in individual recruited neurons, where higher amplitudes resulted in stronger activation.

#### Mean distance of recruited neurons is changed only slightly with biomimetic amplitude modulation

Lastly, given that we found recruitment and intensity changes with dynamic parameter modulation, we also wanted to know if distance was changed to understand if the onset/offset ICMS changes were primarily driven by changes in total neurons recruited near the electrode or if these changes were due to an expanded volume of activation. Therefore, we investigated how the average distance changed across different train types, durations, and amplitudes. With clinically relevant Short trains, there were no significant differences (*p*=0.15, Fig. 3H), although BioBoth trains tended to recruit neurons farthest from the electrode (200±3 µm) and Fixed trains recruited neurons closest to the electrode (180±3 µm), representing a 10% difference in distance.

Next, to understand if distance was changed substantially during each of the phases, we looked at Long trains and found that BioAmp trains evoked activity in neurons farthest from the electrode in onset and offset phases, representing a 13-28% difference (p<0.001, Fig. 3I left,right). Interestingly, there were no significant differences in the distance of neurons recruited during the hold phase between trains (*p*=0.69, Fig. 3I center), despite the large decrease in amplitude for the BioAmp/BioBoth trains. For BioAmp and BioBoth trains, we expected there might be a decrease in the average distance from the electrode during the hold phase compared to the onset, because of the large reduction in amplitude. However, neurons recruited during the hold phase were not significantly closer to the electrode than neurons recruited during onset (*p*=0.28 and *p*=0.11, Fig. 3I), although the mean during the hold phase was closer for BioAmp (214±26 µm) and BioBoth (186±21 µm) than during the onset phase (BioAmp=242±6 µm, BioBoth=223±6 µm), representing a 12-17% difference. For BioAmp trains, more distant neurons were able to recover from DCS during the low hold amplitude which allowed strong offset responses to evoke calcium activity significantly farther from the electrode than Fixed trains (*p*<0.01; Fig. 3I right). Long+ trains had similar trends to Long trains, although the onset phase was not significantly different, indicating that the increased amplitude during the hold resulted in similar distances of recruited neurons across trains (*p*>0.05, Fig. 3J). Together, biomimetic amplitude changes could affect the distance of recruited neurons. However, the differences in average distance of recruited neurons were generally smaller in magnitude (up to 28%) than the differences in recruitment (up to 53%) and mean intensity (up to 46%) across trains and phases, indicating that the population response is dominated by increases in recruitment and mean intensity in neurons closer to the electrode.

Overall, biomimetic amplitude modulation increased the number of recruited neurons, mean intensity, and average distance of neurons recruited during onset and especially during offset. In contrast, BioFreq and Fixed trains increased the number of neurons active during the hold phase which depressed over time with continuous ICMS, resulting in low activity at offset. BioBoth trains had profiles and distributions similar to BioAmp trains, suggesting that amplitude modulation dominated the effects of BioBoth trains. Short BioBoth trains did have some small and significant decreases in mean intensity (Fig. 3E) and small and non-significant decrease in recruitment (Fig. 3A) compared to Short BioAmp trains. These differences could indicate that the high frequency phases in the BioBoth train resulted in decreases in intensity during the onset/offset phases. Long+ trains increased the number of neurons recruited and intensity during the hold phase relative to Long trains, resulting in more DCS (Fig. 3D). For biomimetic amplitude modulated trains, the increased DCS driven by Long+ trains resulted in a non-significant decrease in recruited neurons during the offset relative to Long trains (Fig. 3D). Differences in recruitment and intensity were larger than differences in distance, suggesting differences in network activation are primarily due to changes in the number and intensity of neurons recruited. These findings clarify the mechanisms by which biomimetic amplitude modulation substantially changes the population response and demonstrated that increases in recruitment and intensity during the hold phase (via amplitude modulation) directly affect the excitability of the offset phase.

### Higher and longer amplitude and frequency modulation induces weaker offset responses and stronger depression in some individual neurons

Higher amplitudes and frequencies of ICMS evoke higher intensity calcium activity and induce stronger depression in recruited populations, but previous studies showed that the degree of response and depression is heterogeneous across individual neurons [34], [35], [37], [38]. Having found that the intensity of individual neurons is modulated by dynamic parameter modulation, we were interested to know if dynamic modulation of ICMS amplitude and frequency evoked greater DCS and offset responses in individual neurons. To evaluate the heterogeneity of this effect, GCaMP responses were classified as “Modulated” if an increased offset response was evoked or “Unmodulated” if an offset response was not detected. “Modulated” classes were further divided into three subcategories: higher intensity at onset (Onset), offset (Offset), or no difference (Same). Similarly, “Unmodulated” classes were further subclassified based on how quickly they depressed to ICMS. Because the degree of depression was different for Short and Long trains (due to different durations of ICMS) the criteria we used to identify depression was different across Short and Long/Long+ trains. For Short trains, we defined Rapidly-adapting (Short RA=<85% after 1 s, criteria), Slowly-adapting (Short SA=<95% after 1 s, criteria), or Non-adapting (Short NA>95% after 1 s, criteria).

#### Amplitude and frequency modulation evoke weaker offset responses and stronger depression in some individual neurons for Short trains

We were interested in understanding how modulation of amplitude and frequency in Short trains influenced the distribution of offset calcium activity and DCS, given that Short trains have been shown to have heterogeneous effects on perception in humans [10]. We found that all neurons for BioAmp and BioBoth trains were “Modulated” (Fig. 4A,B, Supp. Fig. 4). The distribution of classes between BioAmp and BioBoth trains were similar except for an increase in the number of “Offset” neurons for BioAmp trains and corresponding increase in “Same” neurons for BioBoth trains. In contrast, BioFreq trains only modulated offset transients in a fraction of the neurons (Modulated, 5%, Fig. 4C). Because of how small the modulated population was, the average response was not impacted (Fig. 2) but at least a small subgroup of neurons were modulated by dynamic changes in frequency. The BioBoth train then seems to be dominated by amplitude effects, so that the evoked response was similar, but not identical to the BioAmp train. However, the frequency modulation in the BioBoth train may have resulted in weaker responses during the Offset (Fig. 4B), which could be why they had lower mean intensities relative to BioAmp trains (Fig. 3E). Lastly, Fixed trains did not evoke any offset transients (no modulation) and had a much greater subpopulation of RA neurons (83%) compared to BioFreq trains (23%) (Fig 4C,D). For Short trains, we found then that both dynamic amplitude and frequency modulation can evoke offset transients. However, frequency modulation had a more limited capacity to modulate an offset response so that BioBoth train responses were similar to those of BioAmp trains. The high frequency during the high amplitude transients of the BioBoth trains may have increased DCS relative to BioAmp trains (resulting in less “Offset” responses), while the low frequency during the high amplitude hold phase for BioFreq trains may have decreased DCS relative to Fixed trains (resulting in less “RA” responses). The interaction between amplitude and frequency for dynamic parameter modulation then may be an important consideration for biomimetic modulation.

**Figure 4:**
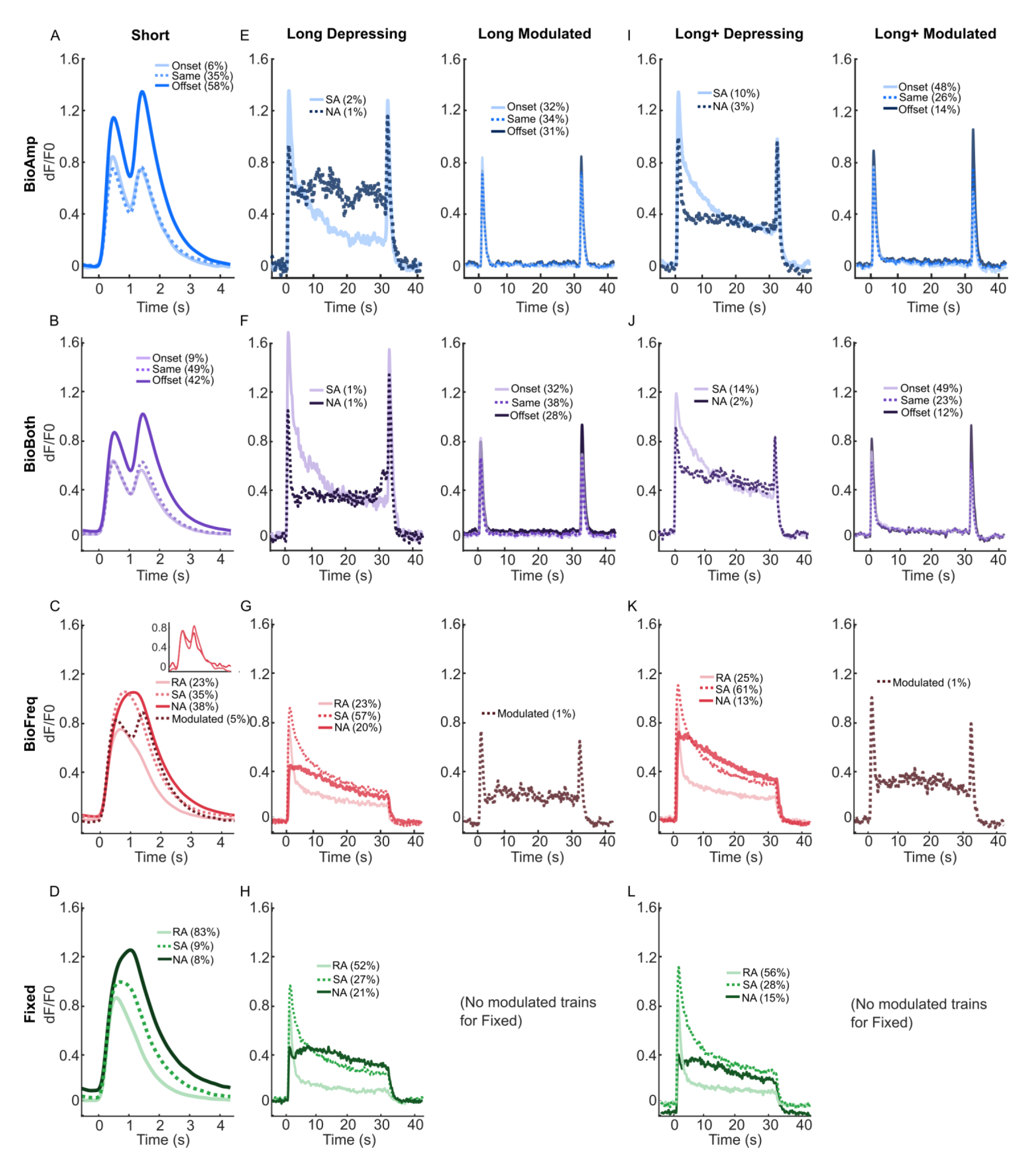
Biomimetic amplitude modulation drives greater activation during hold and offset for some neurons, while biomimetic frequency lowers depression of recruited neurons relative to fixed trains. Each panel shows the corresponding average traces for each class. A-D) Response to Short trains. A-B) Individual neurons have the greatest intensity either at the onset phase, offset phase, or had the same intensities at both peaks for BioAmp (A) and BioBoth (B) trains. C-D) Individual neurons showed either rapid adaptation (<85%, RA), slow adaptation (<95%, SA), or no adaptation (NA) following 1 s of stimulation for BioFreq (C) and Fixed (D) trains. A small subpopulation were also responsive to the onset and offset of ICMS (Modulated) for BioFreq trains. A few of these modulated neurons are highlighted in the inset. E-H) Response to Long trains. Classes for each train are split between two plots to improve visualization of profiles (based on modulation or depression). E-F) Individual neurons respond similarly to the Short trains for BioAmp (E) and BioBoth (F) trains, but also contain some neurons with elevated activity during the hold, which are divided into slowly adapting (SA) and non-adapting (NA) type responses. G-H) Individual neurons can be divided into those that show rapid adaptation in the first 10-s of ICMS (RA), those that have slow adaptation throughout the 30-s of ICMS (SA), or those that have no adaptation throughout (Non-adapting, NA) for the BioFreq (G) and Fixed (H) trains. Similar to Short trains, a small subpopulation of neurons were modulated by BioFreq trains. I-L) Response to Long+ trains. I-J) Individual neurons respond similarly to the Long trains except with more neurons active during the hold phase for the BioAmp (I) and BioBoth (J) trains and more depressing responses across all trains. K-L) Individual neurons were divided into identical classes as the Long trains for BioFreq (K) and Fixed (L) trains.

#### Amplitude and frequency modulation with longer hold periods evoke weaker offset responses compared to onset response with stronger depression than Short trains

Although BioAmp, BioBoth, and BioFreq trains were able to modulate offset calcium transients, if neurons maintain this offset modulation after prolonged DCS from long durations of ICMS was unclear. Therefore, we next investigated how extending the modulation period of amplitude and frequency for 30-s Long trains impacted DCS in individual neurons and if increased DCS reduced the offset response. Neurons that had strong onset and offset transients but low activity during the hold phase were classified as Onset, Offset, or Same. For BioAmp and BioBoth trains, 100% of the neurons demonstrated an offset transient (Fig. 4E,F). In contrast, only 1% of neurons were modulated under BioFreq Long trains, suggesting that 4% out of 5% of the Modulated neurons with BioFreq Short train became depressed during Long trains and were unable to drive an offset response (Fig. 4G, right). As expected, Fixed trains did not modulate any offset activity (Fig. 4H). Increasing stimulation amplitude (Long+) did not decrease the number of Modulated neurons for BioAmp (100%), BioBoth (100%), nor BioFreq (1%) (Fig. 4I-L). DCS evoked by long durations of ICMS reduced the number of Modulated offset responsive neurons for BioFreq, but not BioAmp and BioBoth trains. However, the longer DCS duration did reduce the number of “Offset” type neurons for BioAmp/BioBoth trains, suggesting a reduction in the offset peak driven by the longer duration. The distribution of classes was more similar between BioAmp and BioBoth Long/Long+ trains than they were for Short trains (Fig. 4A,B,E,F,I,J). This may be because the extended hold phase allowed neurons to recover, so that neurons initially depressed by the high-frequency and high-amplitude transient of the BioBoth train were able to recover. Together, Long trains increased depression of neurons relative to Short trains, although the long hold period may have reduced the impact of the onset transient on the offset transient activity.

#### Amplitude modulation induces depression in neurons recruited during the hold phase but reduces number of recruited neurons, while frequency modulation lowers depression of recruited neurons

BioAmp and BioBoth trains modulated 100% of recruited neurons even following long durations of ICMS (Fig. 4E,F,I,J), but only a small percent of neurons maintained activity during the long hold phase (2-3% for Long and 13-16% for Long+ trains). Having found that BioAmp and BioBoth trains seemed to reduce depression by lowering recruitment and intensity, we next wanted to know if neurons that were continuously recruited during the hold phase still experienced DCS. For Long trains, only 3% of BioAmp responses and 2% of BioBoth responses had sustained activity during the hold phase (Fig. 4E,F left), supporting the principle that low amplitude hold period can reduce the number of neurons undergoing DCS. Calcium response profiles of neurons that sustained activity during the hold phase were further divided into SA and NA populations based on the rate of depression (RA type neurons could not be distinguished from neurons with low activity during the hold period).

We next wanted to understand if the rate and magnitude of DCS then affected the magnitude of the offset response. SA neurons had a decrease in intensity for the offset peak relative to the onset peak (avg. decrease peak-to-peak, Long: BioAmp=0.5±7.9%, BioBoth=12.5±7.8%,) demonstrating a decrease in excitability of the recruited neurons (Fig. 4E,F). NA neurons maintained steady calcium activity throughout the hold phase or showed slight facilitation. These NA neurons showed an increase in the offset peak for Long trains (avg. increase peak-to-peak: BioAmp=19±6.3%, BioBoth=31±14%) (Fig. 4E,F), although it is difficult to know if this is an increase in the excitability of the neuron or an artifact of the calcium dissociation latencies contributing to the offset response. Nevertheless, these results indicate that a small percent of neurons respond during the low amplitude hold period for Long trains, some of these sustained activity neurons are susceptible to DCS, and this depression decreases the ability of neurons to respond to the offset transient. These findings support the claim that BioAmp/BioBoth trains are able to maintain excitability of the network by reducing the recruitment of neurons, since a large percent of neurons recruited during the hold phase did show decreases in excitability. BioFreq Long trains had fewer RA neurons (27%) compared to Fixed trains (57%) suggesting that frequency modulation can also reduce DCS, even if there are far fewer neurons that have an offset response. Taken together, amplitude modulation decreases the number of recruited neurons which reduces DCS, whereas frequency modulation decreases DCS of recruited neurons.

#### Increasing amplitude during continuous ICMS results in more depression in individual neurons and corresponding decreases in the offset response

Given that amplitude and frequency modulation can impact DCS and higher amplitude trains recruit more neurons [34], [87], we investigated how increasing ICMS amplitude (Long+ trains) influenced the number of neurons active during the hold period and overall DCS. Increasing ICMS amplitude (Long+) increased the number of Modulated neurons with elevated activity during the hold period for BioAmp (12%) and BioBoth (16%) but did not change the amount of Modulated neurons for BioFreq (1%, Fig. 4I-K). For both BioFreq and Fixed Long+ trains, increasing ICMS amplitude led to an increase in RA neurons (31% vs 21% and 65% vs 57%, respectively) compared to Long trains. Fixed trains evoked more RA responses (Long=52%, Long+=58%) than BioFreq trains (Long=23%, Long+=25%), likely because the higher frequency during the hold phase depressed neurons more strongly and quickly. Long+ trains also evoked less NA responses (Long+: BioFreq=11%, Fixed=12% vs. Long: BioFreq=16%, Fixed=19%) than Long, likely because the higher amplitude depressed neurons more strongly and quickly. The higher amplitude of the hold phase for the Long+ trains caused more neurons to experience DCS, resulting in a decrease in the average offset peak intensity vs. Long trains (avg. decrease peak-to-peak: BioAmp=14±1.0%, BioBoth=16±1.0%). This decrease in offset peak driven by the Long+ trains resulted in much less Offset classified neurons (15 and 12%) compared to the Long trains (31 and 28%). Taken together, higher amplitudes increase the number of neurons active during the hold phase which increases DCS and reduces the offset response.

Overall, we found that dynamic amplitude and frequency modulation influence some individual neurons to reduce DCS and enhance encoding of offset transients. Higher amplitudes, frequencies, and durations of ICMS evoked more DCS, which produced more “Onset” and “RA” type responses. Increasing amplitude also increased the number of neurons activated during the hold period, and many of these neurons had decreased responses to the offset transient, demonstrating a decrease in excitability. The responses evoked by BioBoth trains were similar in shape and distribution to those evoked by BioAmp but were substantially different than the responses evoked by BioFreq trains, implying that the effects of BioBoth trains are dominated by amplitude modulation. However, BioBoth Short trains had much less Offset responses than BioAmp Short trains, which could indicate increased depression driven by the high-amplitude, high-frequency transients. We also demonstrated that the total percent of neurons represented by each class was similar across somatosensory and visual cortices (Supp. Fig. 5), providing further support that the ICMS evoked spatiotemporal activations of somatosensory and visual cortices are similar. Taken together, by categorizing individual neurons with different responses to the same ICMS parameters based primarily on modulation and depression, we found that; 1) different neurons respond heterogeneously to the same parameters for all trains applied here, 2) a small subpopulation of neurons modulate to dynamic frequency changes, and 3) decreasing frequency during high amplitude periods can reduce DCS resulting in more “Offset” and “NA” type responses. These findings clarify that the effects of biomimetic modulation are not identical for all recruited neurons and emphasize the importance of individual neuron responses as well as the interaction between amplitude and frequency for parameter modulation.

**Figure 5:**
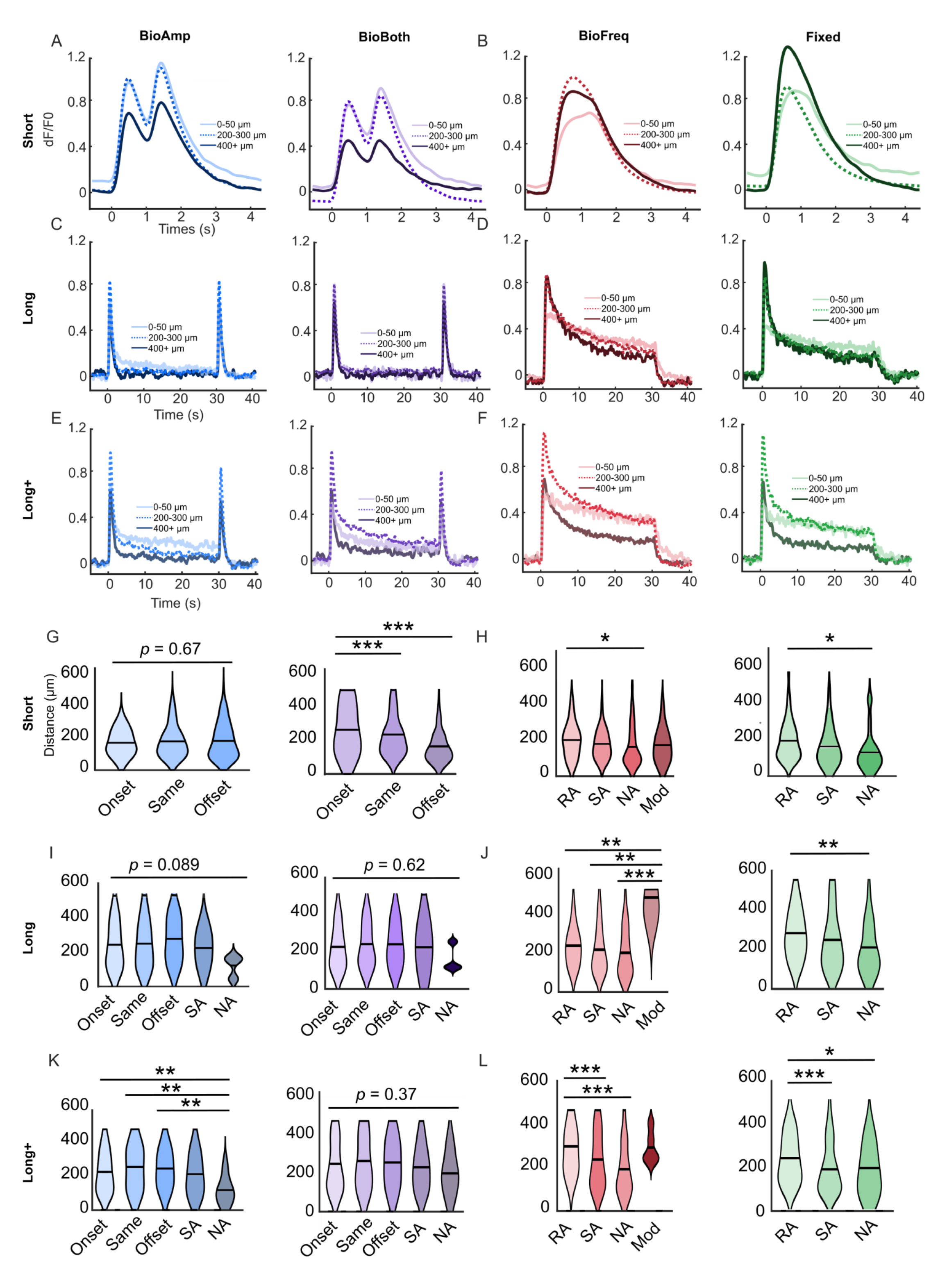
Biomimetic amplitude modulation reduces relationships between distance and depression. A-F) Average neural response to Short (A,B), Long (C,D), and Long+ (E,F) trains based on distance from the electrode. Generally, biomimetic amplitude modulated trains had higher intensity and facilitating responses closer to the electrode with lower intensity and depressing responses farther from the electrode, while BioFreq and Fixed trains had lower intensity and facilitating responses closer to the electrode and higher intensity and depressing responses farther from the electrode. G,H) Violin plots showing the distance of neurons separated by class for Short trains. The solid black line indicates the mean for each class. Neurons farther from the electrode were more likely to be identified as “Onset” or “Rapidly Adapting (RA), indicating depression, while neurons closer to the electrode were more likely to be identified as “Offset” or “Non-adapting” (NA) indicating stability or facilitation to ICMS. I,J) Violin plots showing the distance of neurons separated by class for Long trains. For Long Fixed and BioFreq trains, RA neurons were closer to the electrode and NA neurons were farther, but there were no significant relationships for BioAmp and BioBoth Long trains. Modulated BioFreq responses were also significantly farther from the electrode. K,L) Violin plots showing the distance of neurons separated by class for Long+ trains. BioAmp trains had 3% NA neurons that were active during the hold phase and significantly farther from the electrode, demonstrating that distance depression relationships still exist for the small number of neurons active during the holds period.

### Biomimetic amplitude modulation results in weaker relationships between neural depression and distance from the electrode

Having found in the temporal responses that some neurons experienced greater depression and weaker offset responses with amplitude and frequency modulation, we next focused on investigating the spatial responses to dynamic amplitude and frequency modulation. Previously, neurons farther from the electrode have been found to depress more than neurons close to the electrode [34], [35], [37]. Therefore, we hypothesized that the class of each neuron, and associated DCS, would be related to the distance from the electrode. Here, we found that the evoked intensity and temporal profile were directly related to the neuron’s distance from the electrode (Fig. 5).

#### Amplitude and frequency modulation evoke more “Onset” and “RA” responses in neurons farther from the electrode, and more “Offset” and “NA” responses in neurons closer to the electrode for Short trains

To investigate the role of distance on calcium intensity of activation, we plotted the average calcium profile for three distance bins: 0-50, 200-300, and >400 µm. For Short BioAmp and BioBoth trains, the farthest neurons had the lowest evoked intensities (Fig. 5A), so that neurons >400 µm were significantly less intense than neurons in the 200-300 µm bin in the first 1 s of ICMS (*p*<0.05, Supp Fig. 6I). For Short BioFreq and Fixed trains, differences in distance were not significant (Supp. Fig. 6I) although the nearest neurons had the lowest intensities in the first 1 s of ICMS (Fig. 5B). For all trains, the neurons farthest from the electrode appeared to undergo more depression: 1) for Short BioAmp/BioBoth trains the farthest neurons were more “Onset” while the closest neurons were more “Offset”; and 2) for Biofreq/Fixed trains the farthest neurons were “RA” with decreasing calcium activity over the stimulation period while the closest neurons were more “NA” with increasing calcium activity (Fig. 5A,B). “Onset” and “RA” type neurons seem to represent neurons with decreases in excitability, while “Offset” and “NA” neurons seem to represent neurons that are stable or facilitating to ICMS. Short trains of all types then are driving stronger depression of neurons farther from the electrode, similar to what has been shown previously with 30 s fixed trains [34], [35], [37].

**Figure 6:**
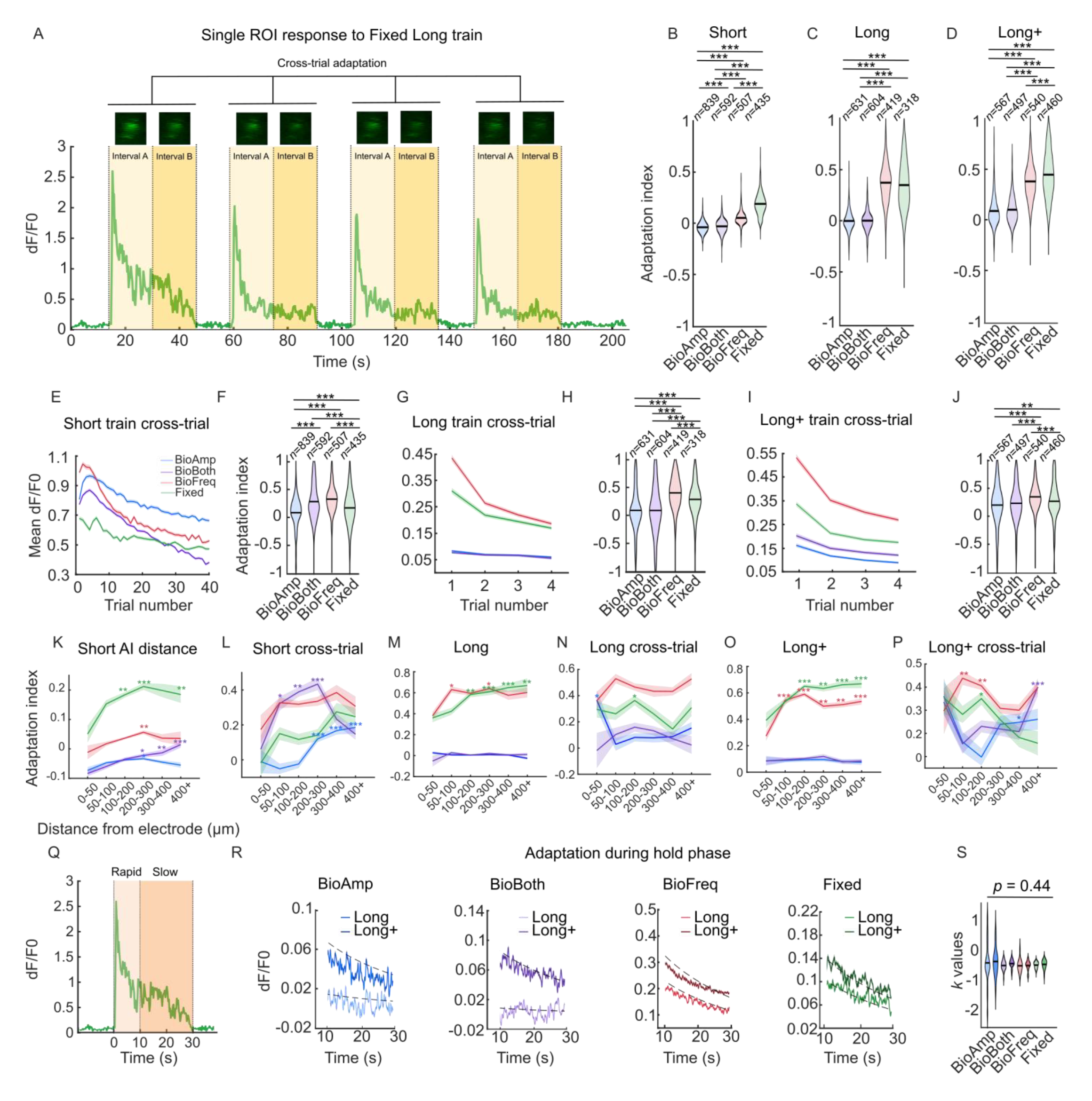
Biomimetic amplitude and frequency modulation can reduce rapid depression relative to fixed parameter stimulation, while slow depression occurs under intensity-dependent mechanisms. A) An example of a single neuron showing depression within trials (DCS) and depression across trials (SIDNE) to a Fixed Long train. Adaptation indices for continuous stimulation were calculated using Intervals A and B with Equation 1. Cross-trial depression was calculated using the average intensity from each trial and using the first and last trial. B-D) Violin plots showing adaptation indices for each Short (B), Long (C), and Long+ (D) trains. Adaptation indices (when comparing the first half to the second half of the train) were greater for the BioFreq and Fixed trains than the BioAmp and BioBoth trains for the Short (B), Long (C), and Long+ (D) trains, indicating stronger depression. BioFreq trains evoked less depression than Fixed trains for Short and Long+ trains, except for Short BioBoth trains. E-J) Biomimetic amplitude modulation generally decreases depression across trials for Short trains (E-F), Long trains (G-H), and Long+ trains (I-J). E,G,I) The mean dF/F0 of all active neurons is plotted across trials and the standard error is marked by the shaded region. F,H,J) Adaptation indices for all active neurons are shown by the violin plot. K-P) Average adaptation indices calculated as a function of distance from the electrode for continuous adaptation (K,M,P) and cross-trial adaptation (L,N,P) for Short (K,L), Long (M.N), and Long+ (O,P) trains. Shaded regions show the standard error across neurons in each distance bin. Asterisks indicate Ais that were significantly larger than the lowest AI for the given train. Q) The same trace from (A) is used to demonstrate the division of the Long and Long+ trials into the “rapid” and “slow” adaptation phases. R) Adaptation during the slow phase occurs consistently as a function of the starting intensity across trains. Average intensity traces for each Long and Long+ train are plotted for the hold phase. Adaptation during the hold period can be accounted for with Equation 2, which is shown by the dotted line. S) Violin plots showing the adaptation slopes for each train during the slow phase (*n*=21-26). *Note that none of these slopes are significantly different*.

#### Long trains evoke more depression in neurons farther from the electrode for Fixed and BioFreq trains with weaker relationships between distance and depression for BioAmp and BioBoth trains

While Short ICMS trains might increase depression of neurons farther from the electrode, how the low intensity hold phase might affect these relationships or how the network might behave during extended bouts of ICMS was not clear. To elucidate the balance of facilitation and depression at equilibrium and how longer hold durations affect the offset response, we next investigated Long trains. Long BioAmp and BioBoth train responses were consistent across distance, with a small increase in the hold activity for neurons closest to the electrode (Fig. 5C). Long BioFreq and Fixed trains evoked depressing responses farther from the electrode, and non-adapting responses closer to the electrode (Fig. 5D). Across all trains, neurons in the 200-300 µm range tended to have the higher mean intensity than >400 µm in the first 1 s of ICMS, although this difference was only significant for the BioAmp trains (*p*<0.001, Supp. Fig. 6J).

We also looked at Long+ trains to better understand how increasing the amplitude during extended hold phases might affect the relationship between depression and distance and the offset response. Trends for Long+ trains were similar to Long trains with neurons closer to the electrode having higher hold activity for the BioAmp/BioBoth trains (Fig. 5E) and non-adapting responses for the BioFreq/Fixed trains, indicating that the relationship between depression and distance is consistent at higher amplitudes despite changes in the overall depression of the network (Fig. 5F). Neurons in the 200-300 µm range still had the highest mean intensity in the first 1 s of ICMS across trains and all trains had significant differences between the 200-300 µm bin and the >400 µm bin (*p*<0.05, Supp. Fig. 6K). The higher amplitude of the Long+ train likely induced more rapid depression in neurons >400 µm, resulting in significantly lower mean intensity in the first 1 s (*p*<0.05). For BioAmp/BioBoth trains, neurons in the 0-50 µm range had more stable responses during the hold period than neurons in the 200-300 µm range, despite both groups showing maintained hold activation (Fig. 5E). These results imply that neurons recruited directly by ICMS (closer to the electrode) can maintain excitability to ICMS despite continuous hold activation while those indirectly recruited undergo DCS. However, this increased stability could also be due to ATP depletion and corresponding inability to remove Ca++ from the cell. These findings provide further support that biomimetic amplitude modulation can reduce depression and encode offset by decreasing recruitment of neurons during the extended hold phase. Together, these results show that neurons farther from the electrode are more likely to undergo depression induced by ICMS, while neurons closer to the electrode are more likely to be stable/facilitating and to be activated by low amplitudes. However, because BioAmp/BioBoth trains reduce recruited neurons during the extended hold phase, they reduce depression across neurons and correspondingly have weaker relationships between depression and distance.

#### Neurons farther from the electrode are more likely to modulate to biomimetic frequency and are classified as more depressing when recruited continuously

Based on the established relationship between depression and distance, we expected that distance would also correlate with our classification for neurons, which seemed to be primarily driven by depression. We found for the Short BioBoth trains, “Onset” neurons were significantly farther from the electrode than “Offset” and “Same” neurons (*p*<0.001, Fig. 5G right), indicating these distant “Onset” neurons were more likely to undergo DCS. Similarly, for Short BioFreq and Fixed trains, “RA” neurons were significantly farther from the electrode than “NA” neurons (*p*<0.05, Fig. 5H), suggesting that distant neurons experienced greater DCS. By breaking down neuron responses into distance bins for BioBoth trains, we find that neurons >400 µm from the electrode were more likely than average to be “Onset” neurons, while neurons <100 µm from the electrode were more likely than average to be “Offset” neurons (Supp. Fig. 6C). Similarly, neurons less than 50 µm from the electrode were much more likely than average to be “NA” neurons for BioFreq and Fixed trains (Supp. Fig. 6B,D). For Long trains, BioAmp and BioBoth had no significant differences in distance between classes (Fig. 5I), but Fixed trains again had “RA” neurons significantly farther from the electrode than “NA” neurons (*p*<0.01, Fig. 5J right). It appeared that the “NA” neurons that remained active for Long BioAmp and BioBoth trains were closer to the electrode (Fig. 5I), but this was not a significant difference likely because of the small number of neurons that remained active during the hold phase (<3%). Similar to Short trains, neurons <50 µm from the electrode were more likely than average to be “NA” for BioFreq and Fixed Long trains (Supp. Fig. 6F,H).

Knowing that increasing the amplitude would increase the number of neurons active during the hold period, we next evaluated how Long+ trains affected relationships between depression and distance. Neurons were more active during the hold period for Long+ BioAmp/BioBoth trains, which resulted in more SA/NA neurons. For the BioAmp Long+ train, the NA neurons were significantly closer to the electrode than the Onset, Same, and Offset neurons (*p<*0.01, Fig. 5K left). This result suggests that relationships between depression and distance from the electrode still exist for BioAmp/BioBoth trains, where non-adapting neurons are closer to the electrode, but in a much smaller population than Fixed/BioFreq trains. SA and NA neurons only made up a small percent of responses to Long and Long+ BioAmp/BioBoth trains (2-16%, Fig 4E,F,I,J, which would indicate 84-98% of neurons recruited by 30 s BioAmp/BioBoth trains do not have significant relationships between distance and depression. BioAmp/BioBoth trains may reduce or abolish depression among most neurons by reducing recruitment during the hold phase, so that most neurons have similar depression of activity regardless of distance. Long+ BioFreq/Fixed trains had similar relationships to Long trains, with RA neurons significantly farther from the electrode than both SA and NA neurons *(p*<0.05, Fig. 5L).

Long BioFreq trains “Modulated” neurons were also significantly farther from the electrode (*p*<0.01) implying that these neurons may be indirectly activated (Fig. 5J left). For Long+ BioFreq trains, “Modulated” neurons were not significantly different from other groups but were the farthest from the electrode on average (mean distance=282 µm vs. 230 µm all other classes, Fig. 5L left). However, there were so few neurons that were modulated by BioFreq Long trains (3 for Long and 8 for Long+) that drawing strong conclusions about the relationship between distance and depression for this group is difficult. Together, we found that neurons farther from the electrode undergo more DCS across Short, Long, and Long+ trains, resulting in more “Onset” and “RA” type responses. For Long+ trains, only NA neurons for BioAmp trains were significantly closer to the electrode than other Modulated classes (*p*<0.01, Fig. 5K left). This may suggest that only neurons that are continuously recruited have significant relationships between depression and distance, likely because continuous recruitment contributes to stronger DCS.

#### Individual neurons that depress more strongly to Short trains also tend to depress more strongly to Long trains

Finally, given the large amount of literature using 1 s trains in humans and NHPs and the large amount of literature using 30 s trains in rodents, we wanted to know if the responses evoked within individual neurons were similar and related between Short and Long trains. Therefore, we investigated if classifications (which were based primarily on depression and modulation) were themselves correlated across Short and Long trains to better understand if neuron responses are consistent across different lengths of ICMS. Indeed, we found that neurons were more likely to be identified as the same class for Short and Long/Long+ trains (Supp. Fig. 7). For example, “RA” neurons for the Short Fixed trains were more likely than average to be identified as “RA” for the Long and Long+ trains, despite different thresholds for classification between trains (Supp. Fig. 7D,H). Individual neuron responses, and explicitly their rates of DCS, then seem to be consistent to specific amplitudes and frequencies, despite different durations of ICMS. These findings indicate that; 1) evoked responses of neurons obtained with Short trains are related to evoked responses obtained with Long trains based on depression and modulation, and 2) neurons respond consistently to dynamic ICMS profiles of different durations based on their distance from the electrode.

**Figure 7:**
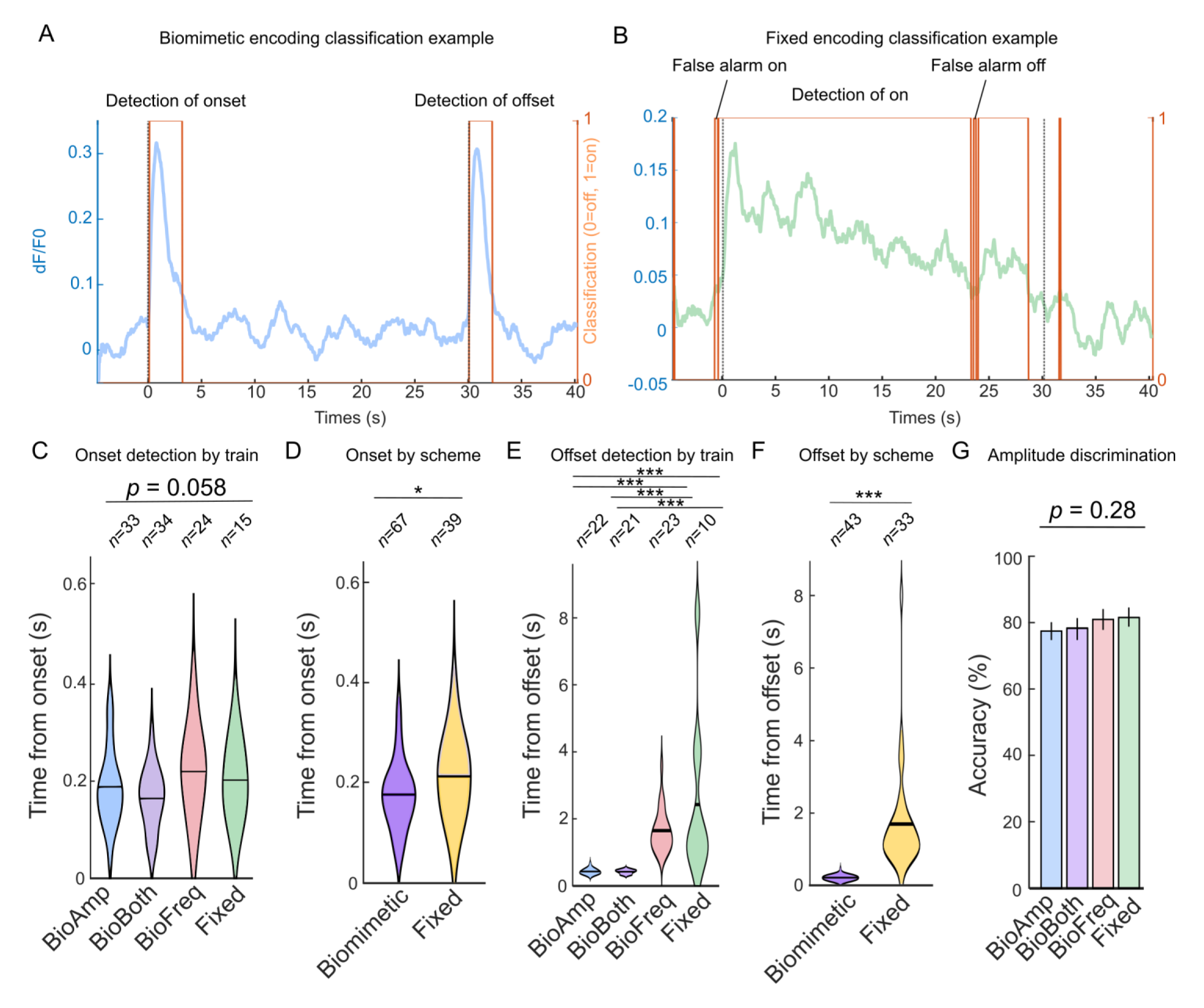
Bayesian classification of ICMS evoked profiles demonstrates improved identification of onset and offset for biomimetic amplitude modulated trains. A) Example of classification of a Long BioAmp train response using the biomimetic encoding model. The blue trace shows the fluorescent intensity evoked by ICMS and the orange trace shows the classification of the onset and offset state (based on confidence>75%). The black dotted line indicates the real stimulation onset and offset. B) Example of classification of a Long Fixed train response using the fixed encoding model, with examples of false alarms for stim on and stim off. A false alarm was defined as confidence of on>75% before stimulation onset or confidence in off>75% before stimulus offset. C-D) Biomimetic encoding with the amplitude modulated trains improved onset detection slightly. C) Violin plot of time to onset detection for each train type. Time to onset was calculated as the time point in which 75% confidence was achieved in onset. Trials on which onset was determined before stimulus onset were considered false alarms and excluded from analysis. The size of the violin represents the kernel density estimate of the *n* trials, the black bar shows the mean. D) Time to onset detection for each encoding scheme. Results from the BioAmp and BioBoth trains were for the biomimetic encoding model while results from the BioFreq and Fixed trains were combined for the Fixed model. E) Biomimetic encoding greatly reduced the time to offset detection. F) Time to offset detection for each encoding scheme. G) All trains resulted in significant accuracy (>75%) at discriminating high amplitude from low amplitudes. Accuracy for amplitude discrimination for each train type across all trains (*n*=28 trials). Total accuracy was based on the correct identification of each time point as being the lower or higher amplitude train. Error bars show standard error.

Given the small number of neurons that were modulated by frequency, we expected that relationships across different durations would be weak. Surprisingly, there were clear relationships for BioFreq “Modulated” neurons between Short and Long trains (Supp. Fig. 7B,F), indicating these neurons represent a unique subclass that could consistently respond to dynamic changes in frequency. However, relationships between Short and Long trains were stronger than relationships between Short and Long+ trains, indicating that the increased amplitude can change individual neural responses, likely by increasing depression. Increased depression may reduce the ability for neurons to modulate to changes in frequency. Long and Long+ trains also evoked similar responses overall, with Long+ trains driving stronger activation during the hold period and more depressing responses (Supp. Fig. 7I). Many “SA/NA” responses for Long+ BioAmp/BioBoth came from “Offset” responses for Long BioAmp/BioBoth trains, because both “SA/NA” and “Offset” responses tended to be closer to the electrode. Together then, these results indicate that individual neurons respond consistently to ICMS trains of different lengths with the same amplitude and frequency, primarily based on their ability to modulate to dynamic changes in parameters and the rate and magnitude of depression.

Overall, we find that the rate and magnitude of depression to ICMS is increased for neurons farther from the electrode, and the depression and modulation of individual neurons were correlated across different durations of ICMS. BioAmp/BioBoth Long and Long+ trains had much weaker relationships between distance and depression than BioFreq/Fixed trains, because the decreased amplitude during the hold phase reduced depression of recruited neurons, so that 84-98% of neurons were not continuously recruited and thus maintained excitability. Long+ trains increased the number of neurons with significant relationships for BioAmp/BioBoth trains by increasing the number of neurons continuously recruited. Neurons that were modulated by BioFreq Long and Long+ trains also tended to be farther from the electrode, implying that they were indirectly activated. Synaptic integration then might be required for encoding of dynamic frequency changes. These findings establish that the depression and modulation of neurons are dependent on their distance from the electrode, but biomimetic amplitude modulation can make responses less heterogeneous and dependent on distance by reducing depression across recruited neurons.

### Biomimetic amplitude and frequency modulation reduces depression of neural calcium activity and excitability during repeated stimulation

We observed in the previous sections that depression of neuron activity is related to the neuron’s distance from the electrode, the length of stimulation, and the modulation of amplitude and frequency. Adaptation, or decreases in perceived intensity, has been previously demonstrated for ICMS in humans [15], [20], where stimulation over relatively short periods could cause sensation to become imperceptible. While sensory adaptation is normal [26]–[30], stimulation induced depression of neuronal excitability (SIDNE) seems to reflect abnormal processes that can take multiple days for recovery [32], [33], and maintaining excitability of neurons is desirable for clinical applications. We hypothesized that biomimetic modulation, by decreasing the amplitude and frequency during the hold phase, could better maintain the excitability of neurons over time. We were able to measure depression during continuous stimulation (DCS) and across trials (SIDNE) in individual neurons (Fig. 6A).

#### Biomimetic amplitude and frequency modulation reduces depression to continuous stimulation within trials

To understand how parameter modulation impacts DCS, adaptation was assessed within trials to continuous ICMS. For continuous ICMS, BioAmp and BioBoth Short trains drove a negative adaptation-index (AI) or minor facilitation (*AI*=-0.04±0.002 and −0.03±0.003, respectively, Eq. 2), while BioFreq and Fixed trains drove depression, although BioFreq trains resulted in less depression than Fixed trains (*AI*=0.05±0.003 and 0.18±0.006, *p*<0.001, Fig. 6B). For Short BioAmp and BioBoth trains, the 2.5 s onset calcium activity and slow calcium dissociation constant of calcium could contribute to the offset response which made it unclear if there was real “facilitation” of neuronal activity (Fig. 4A,B). To clarify differences between onset and offset, we next investigated DCS with Long trains where onset and offset transients were separated by 30 s. BioAmp and BioBoth Long trains drove minor depression (*AI*=0.01±0.004 and 0.012±0.004), while BioFreq and Fixed drove stronger depression (*AI*=0.40±0.009 and 0.37±0.013, *p*<0.001, Fig 6C). Minor depression then occurs with BioAmp and BioBoth Long trains, but not with the Short trains, which could be because calcium activity latencies cause the onset transient to influence the offset transient or because the shorter duration of ICMS allows for facilitation. Because higher amplitudes increase recruitment and corresponding depression, we wanted to see if these relationships disappeared for Long+ trains. Long+ trains increased depression for BioAmp/BioBoth trains (*AI*=0.08±0.007 and 0.1±0.006), which still drove less depression than Long+ BioFreq/Fixed trains (*AI*=0.38±0.009 and 0.42±0.01, *p*<0.001, Fig. 6D). BioFreq Long+ trains drove less depression than Fixed Long+ trains (Fig. 6D, *p<*0.001), although there were no significant differences for Long trains (Fig. 6C, *p>*0.05) indicating that the increased amplitude results in stronger depression for trains with higher frequencies, emphasizing the importance of amplitude and frequency interactions. Together, these results demonstrated that biomimetic amplitude modulation substantially reduces DCS by reducing recruitment during the hold period. However, biomimetic frequency also reduced DCS less substantially relative to Fixed trains, by reducing the frequency of activation of continuously recruited neurons. Amplitude modulation was then able to strongly reduce depression, while frequency modulation more subtly reduced depression for the clinically relevant parameter ranges used here.

#### Biomimetic amplitude modulation reduces depression across trials, while biomimetic frequency modulation increases depression across trials

Continuous ICMS allowed for assessment of depression during ICMS, but we were also interested in how ICMS changed excitability following ICMS cessation. To understand how parameter modulation impacts SIDNE, adaptation was also assessed across trials. Short trains drove facilitation across the first three trials, followed by depression (Fig. 6E). This could again represent real facilitation of recruited neurons with short stimulus trains, but also could be related to overlap of calcium evoked activity. BioAmp Short trains drove the least cross-trial depression (*AI*=0.09±0.01, *p*<0.001, Fig. 6F). Interestingly, BioFreq Short trains initially drove the most intense calcium activity, but then depressed the fastest after 5 Short trains (*AI*=0.33±0.01, *p*<0.001, Fig. 6F). Surprisingly, we found that BioBoth Short trains induced significantly more cross-trial depression than either BioAmp or Fixed trains (*p*<0.001, Fig. 6F). BioBoth trains induced less DCS than BioFreq trains, but did not have significant differences in cross-trial adaptation, which could imply that the frequency component is more important than the amplitude component for cross-trial adaptation for Short trains. Given this surprising result, we were interested to know if trains of longer durations also demonstrated differences between BioAmp and BioBoth trains. For Long trains, no facilitation occurred and the strongest depression occurred from the first to the second trial (Fig. 6G). BioAmp and BioBoth Long trains drove less depression across trials (*AI*=0.1±0.015 and 0.1±0.018, Fig. 6H) than BioFreq and Fixed Long trains (*AI*=0.41±0.014 and 0.29±0.017, *p*<0.001, Fig. 6H). Unlike the Short trains then, cross-trial adaptation for Long/Long+ trains were dominated by amplitude effects, similar to DCS.

We wanted to know if increasing the amplitude changes these relationships given the increased recruitment and intensity of recruitment. For Long+ trains, BioFreq trains had the most cross-trial depression (*AI*=0.34±0.01, Fig. 6J) while BioAmp trains had the least (*AI*=0.21±0.016, *p*<0.001, Fig. 6I,J). Long and Long+ Fixed trains had significantly less cross-trial depression than BioFreq trains (*p*<0.001, Fig. 6G-J), which was the opposite of what we saw for continuous ICMS. While decreased frequencies intuitively would reduce depression for continuous ICMS, why BioFreq trains increased depression across trials is unclear. However, BioFreq trains evoked the most intense responses across trials (Fig. 6G,I). In fact, trains that evoked the least intense responses overall also induced the least cross-trial depression. SIDNE then may be dependent on the overall intensity of recruitment, while DCS is dependent on other mechanisms of depression unrelated to overall intensity. Together, these results demonstrate that biomimetic amplitude modulation reduces SIDNE, likely by reducing the overall intensity of recruitment of neurons, which is increased for biomimetic frequency modulation.

#### Biomimetic amplitude modulation decreases DCS but not SIDNE of neurons farther from the electrode

We also found that adaptation indices were related to distance from the electrode, as predicted from the previous results (Fig. 5). Neurons farther from the electrode for BioBoth, BioFreq, and Fixed Short trains (Fig. 6K), and Fixed/BioFreq Long/Long+ trains (Fig. 6M,O) depressed more to continuous stimulation (*p*<0.05). BioAmp/BioBoth Long/Long+ trains did not have significant relationships between distance and depression (*p*>0.05, Fig. 6M,O). Interestingly, BioBoth but not BioAmp Short trains had stronger DCS for farther neurons (Fig. 6K). These differences provide further support that frequency modulation may play a stronger role in depression induced within Short trains.

For SIDNE (i.e. cross-trial adaptation), there was also generally more depression farther from the electrode for Short trains, although only BioAmp and BioBoth trains had significant differences (*p*<0.05, Fig. 6L). Additionally, BioBoth trains drove stronger depression in neurons from 50-300 µm (*p*<0.05, Fig. 6L), while BioAmp trains drove stronger depression in neurons from 200-540 µm (*p*<0.05, Fig. 6L), highlighting further differences between these two trains. For Long and Long+ trains, SIDNE did not seem to correlate strongly with distance from the electrode (Fig. 6N,P). Differences in the distance relationship between DCS and SIDNE imply that DCS operates under different mechanisms than SIDNE. By reducing recruitment of neurons, BioAmp/BioBoth trains do not generally drive more depression in neurons farther from the electrode as BioFreq/Fixed trains do. However, BioBoth Short trains also induced greater depression both within and across trials than BioAmp trains, implying that frequency may play a role in depression to short bouts of ICMS.

#### Amplitude and frequency modulation drive depression on both rapid and slow timescales, where slow timescales are only dependent on the starting intensity of activation

We observed that most DCS occurred within the first 10 s of ICMS for Long and Long+ trains. Given previous literature indicating multiple mechanisms for different rates of adaptation, we wanted to know if these two phases occurred under different mechanisms [31]. Therefore, we divided DCS into two phases, an initial rapid depression (the first 10 s) and a slower depression (the last 20 s) (Fig. 6Q). Rapid depression (first 10 s) was dependent on the parameters of ICMS delivered (and affected the neuron’s class), where increasing amplitude and frequency both seemed to result in more rapid depression (and thus more “RA” and “Onset” responses, Fig. 4). In contrast, slow depression (last 20 s) was consistent across all parameters used here, and could be described with a simple equation using only the evoked fluorescent intensity at the beginning of the hold phase:

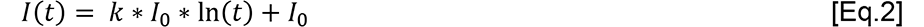

where I(t) is the intensity at time point t, ln(t) is the natural log of the time axis (with range 10-30 s), I_0_ is the intensity at the start of the hold phase, and *k* is a constant where *k*\**I_0_* describes the slope. For all Long and Long+ trains, *k* = −0.43±0.04 was used to fit average traces (Fig. 6R) and *k* values were not significantly different between trains (*p*=0.44, Fig. 6S) indicating consistency in slow depression for all parameters used here. Together, these results indicate that there are two distinct phases of DCS for long periods of ICMS: 1) the first in which higher amplitude and frequencies induce stronger and faster depression; and 2) the second in which depression is dependent on the intensity of recruitment and happens over long periods of time.

Overall, biomimetic amplitude modulation reduced depression within (DCS) and across trials (SIDNE), although depression during the slow depression phase (20-30 s) seemed to be independent of stimulus parameters and only dependent on the starting intensity. BioFreq also reduced DCS relative to Fixed trains (although less dramatically than BioAmp/BioBoth), indicating that lower frequencies can better maintain excitability despite not dynamically modulating most neurons. Some small differences between BioAmp and BioBoth Short trains were found, indicating that frequency may also be more important for depression to Short trains and not completely dominated by amplitude effects. Even though BioFreq trains evoked less DCS, they evoked more SIDNE than Fixed trains, which seems to be due to the higher intensity of recruitment. Additionally, DCS but not SIDNE was dependent on the distance of the recruited neuron from the electrode. Together, these results imply that there are multiple mechanisms under which depression of neural activity occurs with both rapid and slow timescales, and changes in amplitude and frequency have different impacts on these forms of depression. While amplitude had the most dramatic reductions in depression across all forms, frequency is also an important parameter to consider, especially for shorter trains and in interaction with amplitude.

### Ideal observers classify stimulus onset and offset faster with biomimetic amplitude modulation

We found earlier that biomimetic amplitude modulation could result in faster times to peak, which could indicate faster stimulus representation. Previously, increases in stimulus amplitude (which is known to increase neural recruitment and intensity [36], [77], [87], [88]) also resulted in better detection, faster reactions, and higher intensities of ICMS evoked percepts [7], [10], [68], [69], [89]. Therefore, high amplitude transients that signal stimulus onset and offset might lead to more salient percept and improve detection of stimulus onset and offset.

Ideal observer analysis (IDA i.e. Bayesian classification) was used to quantify the strength and speed of neural calcium activity evoked by each train (Fig. 7). Two encoding schemes were used: biomimetic (detection of onset and offset transients) and linear (detection of overall elevation in activity, Fig. 7A,B, Supp. Fig. 8A). BioAmp and BioBoth trains worked best with biomimetic encoding while BioFreq and Fixed worked best with linear encoding (Supp Fig. 8B).

#### Biomimetic amplitude modulation reduces detection times of onset and offset of ICMS for an ideal observer

Biomimetic encoding with BioAmp and BioBoth trains resulted in faster detection of onset (mean detection-time (DT)=0.19±0.014 s and 0.16±0.011 s) than linear encoding with BioFreq and Fixed trains (DT=0.22±0.019 s and 0.20±0.018 s). Although onset times were not significantly different across all trains (*p*=0.058, Fig. 7C), when combined within encoding schemes biomimetic encoding was significantly faster than fixed encoding (i.e. BioAmp and BioBoth vs. Fixed and BioFreq) (*p*=0.019, Fig. 7D). The biomimetic encoding was also much faster at determining offset (DT=0.21±0.016 s and 0.19±0.014 s) compared to linear encoding (DT=1.42±0.13 s and 2.32±0.83 s, *p*<0.001, Fig. 7E,F). Biomimetic encoding also resulted in less false alarms (FAs, early detection) for onset (FAs = 3 and 2) than BioFreq and Fixed (FAs = 12 and 21). Using a simple threshold of +3 baseline standard deviations as a cut-off for detection resulted similarly in better performance for BioAmp and BioBoth trains (Supp. Fig. 7C,D,E). The simple threshold method resulted in less FAs (12 and 17 less) for the onset for BioFreq and Fixed trains than the IDA with increased DTs (0.037±0.012 s and 0.041±0.015 s more, *p*<0.05 paired-sample t-test), but resulted in an increase in FAs (12, 4, 17, and 9 more) for all trains for offset detection (Supp. Fig. 7F,G). Together, these results indicate that for an ideal observer, the strong transients evoked by biomimetic amplitude modulation allow for better classification of the onset and offset of input.

#### Increases in amplitude result in distinguishable activation profiles across all trains for an ideal observer

Intensity of ICMS can be used to determine features of sensory input, such as object compliance [41], [60], and so we were interested in knowing if an ideal observer could distinguish intensities across the different trains used here. Classifiers were additionally trained to discriminate low and high amplitudes (Long vs. Long+). Classifiers were able to discriminate adjusted Long and Long+ fluorescent profiles significantly above chance levels (*p*<0.01, Fisher’s test) for all trains (means=76.9±2.5%, 77.2±3.2%, 82.9±3.2%, and 83.2±3.3%) and these accuracies were not significantly different across trains (*p*=0.28, Fig. 7G). Together, these results indicate that an ideal observer discriminates ICMS intensity based on the evoked activity with significant accuracy for all train types. Overall, biomimetic amplitude modulation improved classification of onset and offset by evoking distinct onset and offset transients and maintained the ability to discriminate different intensities of ICMS.

## Discussion

### Biomimetic amplitude modulation evokes much stronger onset and offset responses compared to biomimetic frequency and fixed modulation

Biomimetic modulation of ICMS may improve the function and quality of sensory feedback [53], [54]. We wanted to know how biomimetic modulation of ICMS affects cortical neuron recruitment. Dynamic changes in amplitude modulated 100% of neurons, while dynamic frequency modulation only modulated a small fraction of neurons (<5%, Fig. 4C). Here, all Short trains were charge matched within clinically relevant parameters, so that differences in total charge injection would not influence the outcome. Comodulation of frequency and amplitude was expected to have the strongest effect on neural calcium activity, since both population size and spike rate have been previously demonstrated to comodulate during high intensity transients [43], [46]. However, frequency of ICMS did not contribute strongly to the onset and offset response in our results and the response to BioBoth trains was dominated by amplitude effects. BioBoth trains only seemed to drive significantly different responses for Short trains, where BioAmp trains evoked higher mean intensity and reduced DCS/SIDNE relative to BioBoth trains (Fig. 3,6). Why BioAmp trains increase mean intensity and decrease depression relative to BioBoth trains is unclear, but may be due to the interaction between amplitude and frequency. In other words, the high amplitude and frequency transients of BioBoth trains induce more DCS for Short trains due to the short break between transients, which also results in lower mean intensity. Indeed, BioAmp trains did evoke more NA type responses than BioBoth trains (Fig. 4A,B) although we cannot be certain this was real facilitation and not due to calcium latencies.

Comodulation for peripheral nerve stimulation has had the strongest effects on perception [58]. However, frequency of ICMS does not behave in the same way as frequency of peripheral nerve stimulation [10]. Microstimulation frequency in the cortex seems to have more complex and electrode specific effects. Here, a small group of neurons (1-5%) were modulated by dynamic frequency changes (Fig. 4C,G,K). The presence of these neurons may affect the local response to ICMS frequency, which could explain different evoked percepts of varying discriminability across electrodes in humans [10], [19]. However, while we found that most neurons did not express clear onset/offset activation with dynamic changes in frequency, frequency is still an important parameter for ICMS modulation, but frequency modulation in high frequency ranges cannot be used to consistently modulate cortical populations to the onset/offset of input. Indeed, neurons can have preferential responses to ICMS frequencies related to distance from the electrode and frequency has strong effects on the intensity of activation and DCS [34], [35]. Lower frequency ranges could likely modulate neurons more strongly, but lower frequencies also result in decreased activation at low amplitudes and have been shown to have highly variable effects on perception in humans [10]. Further studies to understand why different neurons have preferred temporal profiles and with larger parameter sets can shed light on the effects of frequency modulation and how frequency can potentially be modulated in the future to produce more naturalistic sensations.

The inability for BioFreq trains to modulate onset/offset transients is perhaps not surprising in these frequency ranges. Most excitatory neurons fire at low frequencies (average ∼8 Hz), which may indicate that most excitatory neurons are unable to entrain to high ICMS frequencies [74]–[76]. The neurons that were modulated here might represent rare subpopulations of neurons within the cortex which can modulate to much higher frequencies (up to 800 Hz) [93]. More likely, these observed neurons were modulated by complex synaptic integration from other upstream excitatory neurons as well as unlabeled inhibitory neurons with fast firing kinetics, such as parvalbumin-type inhibitory neurons [94]–[97]. Long trains, which were not charge matched, were used to understand how recruitment of neurons is affected by the applied parameters over extended periods of recruitment when some excitatory and inhibitory balance is achieved. We found that neurons farther from the electrode depressed more rapidly, which could be due to recruited inhibition. Interestingly, we found here that BioFreq “Modulated” neurons were significantly farther from the electrode than other classifications for Long trains, implying that these neurons are recruited and modulated indirectly. BioFreq modulated neurons occurring farther from the electrode supports the idea that recruited inhibitory neurons and synaptic integration are required for high frequency modulation of individual neurons. Future work should investigate how amplitude and frequency modulation affect inhibition/disinhibition of excitatory and inhibitory neurons. While the mechanisms underlying the encoding of the dynamic frequency modulation are unclear, in these clinically relevant ranges biomimetic frequency modulation is not consistent in modulating cortical neurons as observed by GCaMP. We found that biomimetic amplitude modulation consistently evoked distinct onset and offset transients, so biomimetic amplitude modulation can consistently evoke neural transients similar to those present in recorded neural data [43]–[52].

### Biomimetic amplitude and frequency modulation maintain neural excitability over long periods of stimulation

Sensory adaptation is a normal part of sensory processing [26]–[31] and may contribute to an increased ability to respond to changes in sensory input. Adaptation of perceived intensity has been observed in peripheral nerve stimulation [98], [99], visual cortex ICMS [15], and somatosensory cortex ICMS [20]. Additionally, stimulation induced depression during continuous stimulation (DCS) and stimulation induced depression of neuronal excitability (SIDNE) have been observed in multiple animal models [32]–[35]. However, how SIDNE or DCS compares to sensory adaptation is not clear. Continuous ICMS in somatosensory cortex can lead to complete loss of perception of ICMS within one minute in humans [20], which could make ICMS ineffective when provided over clinically meaningful stimulation durations. For vibrotactile perception, continuous sensory input results in decreases in intensity over time [26], [28]. However, continuous sensory perception intensities never completely extinguish or fall below zero, and the onset intensities recover on the order of seconds to minutes. In contrast, SIDNE can take multiple days to recover and is accentuated by increases in ICMS amplitude, which seems to indicate a fundamental difference between adaptation evoked by normal sensory input and depression driven by electrical stimulation. Here, biomimetic amplitude modulation better maintained excitability over time for both continuous stimulation (DCS) and across trials (SIDNE) by transiently encoding sensory input.

Biomimetic amplitude modulation reduced the number of neurons recruited during the hold period, which resulted in less depression both within and across trials (Fig. 6B-J). Neurons continuously recruited during the hold phase for BioAmp/BioBoth train still experienced DCS (Fig. 4E,F,I,J). Additionally, this DCS resulted in a reduction in the size of the offset peak, indicating that DCS results in a reduction in excitability of the neuron (Fig. 4E,F,I,J). This reduction in excitability indicates that BioAmp/BioBoth can reduce DCS across the population by decreasing the recruitment of many neurons, where recruited neurons are still subject to depression. However, what exactly is driving depression in recruited neurons remains unclear. DCS could be driven by exhaustion of the neuron, so reduction in recruitment prevents exhaustion. DCS could also be driven by recruited inhibitory drive in the network. Interestingly, neurons that do not have substantial DCS during the hold phase can still have a reduction in the second peak when stimulating at higher amplitudes (Long+, Fig. 4I-K). The reduced excitability then may not be driven by exhaustion of the recruited neurons (since these neurons do not depress during the hold period), which might suggest that reductions in excitability are related to increased inhibitory drive rather than exhaustion. However, reduction in ATP availability induced by ICMS may limit the ability of neurons to pump out Ca++, which results in stable Ca++ during ICMS despite decreases in excitability. To resolve these two possibilities, additional experiments with other modalities (such as electrophysiology, voltage indicators, and/or ATP measurements) are needed. Additionally, SA vs. NA neurons for Long+ trains primarily differ in the first 10 s of ICMS, while in the last 20 s, the average response is overlapping and the offset peaks are similar in size (Fig. 4I,J). The overlapping activity could indicate a balancing of excitation and inhibition is achieved for SA neurons within the first 10 s, so that activation becomes similar to NA neurons. Biomimetic amplitude modulation then may maintain excitability by reducing inhibitory drive, which could represent “disinhibition” of the network (by removing the inhibition that would otherwise be increased during the first 10 s of ICMS). Future work should investigate how dynamic parameter modulation affects the response of both excitatory and inhibitory neurons to shed light onto the mechanisms underlying ICMS induced depression. Here, we showed that biomimetic amplitude modulation decreases depression by decreasing recruitment of neurons during the extended hold phase.

In contrast, biomimetic frequency modulation did not decrease recruitment of neurons during the extended hold phase (Fig. 2,3). However, we still did find that frequency modulation could reduce DCS relative to Fixed trains (Fig. 6). Generally, higher frequencies of electrical stimulation result in stronger DCS and SIDNE for both cortical and peripheral stimulation [20], [34]–[37], [98], [100]. Here, the Fixed train (frequency of 150 Hz) evoked stronger and more rapid depression across 30 s than the BioFreq train (50 Hz), agreeing with previous results [20]. However, the BioAmp train (hold frequency of 150 Hz) did not drive stronger depression than the BioBoth trains (hold frequency of 50 Hz) indicating that higher frequencies alone are not sufficient to increase depression and rather there is an interaction between amplitude, frequency, and duration that is important in depression of neural activity. Specifically, low frequencies at high amplitudes can reduce depression (BioFreq vs. Fixed trains and previous work), but at sufficiently low amplitudes, frequency may not impact the response (BioAmp vs. BioBoth trains). Additionally, Short BioBoth trains induced more DCS than Short BioAmp trains, so that high-amplitude with high-frequencies during the short onset/offset transients may induce more depression, providing further evidence that the interaction between amplitude and frequency is important for depression. Similarly, amplitude and frequency will both have larger effects on depression over extended periods of time (Long and Long+ trains). Therefore, ICMS amplitude, frequency, and duration all likely interact for depression, so that comodulating parameters may have different effects on neural activation and perception i.e. the sum is different than the parts.

Interestingly, BioFreq trains resulted in more SIDNE than Fixed trains (i.e. depression across trials) for all lengths (Short, Long, and Long+) even though BioFreq trains generally resulted in less DCS. BioFreq trains also always resulted in higher intensities of activation than Fixed trains for all durations (Fig 6E,G,I). The higher intensities of activation evoked by BioFreq may have resulted in more SIDNE. For the Long and Long+ trains, BioAmp and BioBoth also result in substantially less SIDNE and lower overall intensity than BioFreq trains (Fig 6E-J).

Therefore, higher overall intensities correlate with stronger SIDNE, regardless of parameters. We also found that depression that occurred within trials after 10 s was directly related to the evoked intensity (Eq. 2). Depression during continuous input in the first 10 s (which is a function of distance from the electrode) may be due to emergent inhibitory recruitment (which may be stronger at high amplitudes and frequencies), but depression that occurs more slowly (after 10 s) and across trials may be related to other metabolic effects or exhaustion of the recruited neurons which are dependent on the intensity of activation (independent of ICMS parameters). Adaptation over short and long periods of time may occur under different mechanisms [28], [39]. Electrical stimulation then seems to evoke at least 2-3 kinds of depression of neuronal activity (rapid DCS vs. slow DCS and SIDNE), which also likely have different mechanisms and times to recovery (where rapid DCS recovers faster). Biomimetic amplitude and frequency modulation then both reduced depression under different mechanisms. More specifically, biomimetic amplitude modulation had the strongest reductions in depression and did so both within and across trials by decreasing recruitment of neurons during the long hold phase. In contrast, biomimetic frequency modulation decreased depression of neurons for continuous stimulation, but increased depression across trials by increasing intensity of activation within trials.

### Mechanisms underlying neural responses to biomimetic ICMS

DCS may occur at two time-scales: rapid and slow [31], similar to the phases we observed here (Fig. 6Q). Rapid and slow DCS may occur under different mechanisms and may depend on intrinsic properties of different cell types and their synaptic connections and role in larger circuits [39]. One possibility is that rapid DCS is driven by recruited inhibition (where higher amplitudes and frequencies of ICMS increased inhibition and corresponding depression) and slow DCS is driven by slower metabolic changes, glia, or plasticity (where higher intensities of activation drive stronger depression) [101], [102]. Indeed, inhibitory neurons play an important role in normal sensory adaptation and different inhibitory subtypes can play different and opposing roles in adaptation [103]– [106]. Specifically, parvalbumin (PV) type neurons (∼6-15% of all neurons) are associated with more general inhibition at higher frequencies, while somatostatin (SOM, ∼6-9% of neurons) type neurons are associated with stimulus specific inhibitions at lower frequencies [95], [97]. ICMS can drive sparse and distributed activation of excitatory neurons by recruiting passing axons [77], [87], [107] but the recruitment of inhibitory neurons may contribute to more focal activation of functional units for feature encoding after onset [108]–[111]. While inhibitory neurons are more sparse than excitatory neurons, ICMS recruits thousands of neurons which likely results in significant inhibitory recruitment. The probability of recruiting different inhibitory subtypes should increase with ICMS amplitude as this increases the density of recruitment, so that higher amplitudes are more likely to recruit different types of neurons.

Here, higher amplitudes and frequencies drove more rapid DCS of neurons, agreeing with previous results [34]– [37], [112]. Increasing amplitude drives DCS in greater number of neurons, while increasing frequency drives faster DCS. Higher amplitudes then may result in increased recruitment of sparsely distributed inhibitory neurons at onset, and higher frequencies drive more rapid DCS of neurons. Indeed, we found that DCS was strongest for neurons farther from the electrode, which agree with previous studies that inhibitory neuron recruitment is increased farther from the electrode [79]. Interestingly, as we increase the amplitude (Long+ train) we see an increase in the rapid DCS of neurons (Fig. 4). In fact, previous work has shown that increases in excitatory neuron activity leads to supralinear increases in inhibitory activation [113]. We also see that for higher amplitudes, the offset peak evoked by BioAmp and BioBoth trains decreases (Fig. 3), which supports the idea that increases in amplitude decreases the excitability of neurons. However, when looking at individual neuron classes, the decrease in the offset peak occurs in SA type neurons and not NA type neurons, except at higher amplitudes (Fig. 4E,F,I,J). NA classified neurons may not be impacted if they are directly antidromically activated. Decreases in excitability of SA neurons then are not driven by the intensity-based DCS that occurs during the hold phase (10-30 s) but are driven by the rapid DCS that occurs in the beginning of the pulse train (0-10 s), which is more likely driven by inhibitory effects. Biomimetic amplitude modulation can reduce DCS then not simply by reducing the intensity-driven depression induced by continuous recruitment, but by reducing inhibitory drive recruited within the first few seconds of ICMS, which most strongly affects neurons farther from the electrode. BioBoth trains also reduce depression in this way, but the high-amplitude high-frequency transients may actually result in more inhibitory recruitment which increased depression for Short trains (Fig. 4A,B). By decreasing inhibitory drive on the recruited population, biomimetic amplitude modulation can maintain the excitability for encoding of the offset of input.

We also find that neurons near the electrode are stable or even slightly facilitated during continuous ICMS with all parameters. These neurons likely represent neurons that are directly recruited by ICMS. Inhibitory input to these neurons would lower the potential for normal firing (through integration of depolarizing inputs), but because electrical stimulation preferentially activates the axon, these neurons may be able to continue to fire despite inhibitory input. However, we are recording Ca++ activity and not spikes directly. Continuous recruitment by ICMS may increase intrinsic cation levels and neurons are unable to achieve rebalancing of ions due to ATP depletion. In this scenario, the increased Ca++ levels would not directly represent firing of the neuron as Ca++ could accumulate in the extracellular space due to insufficient ATP to pump Ca++ ions out of the cytosolic space into the extracellular space or endoplasmic reticulum. Furthermore, firing does not necessarily reflect neurotransmission, and signaling of stimulus evoked response is likely critical for perception. However, previous work has demonstrated elevated glutamate levels near the electrode across the full 30 s of ICMS with high frequencies, suggesting that ICMS does drive continuous neurotransmission even over these longer durations [34]. Although astrocytes reduce glutamate reuptake during high frequency neural activity, so increased glutamate may also not represent continuous release of neurotransmitter [114]–[118]. These possibilities should be explored in the future through detailed investigations combining electrophysiological recordings, calcium imaging, and ICMS.

Most excitatory neurons are not modulated by dynamic changes in frequency within the clinically relevant ranges used here and changes in frequency did not decrease total recruitment, unlike amplitude. Therefore, lowering the frequency likely does not reduce the recruitment of inhibitory neurons during the hold phase. Increasing frequency during the hold phase may increase activity in inhibitory neurons, particularly fast-spiking parvalbumin (PV) type neurons which generally have higher firing rates and may be able to entrain to high frequencies [94]– [96], [119]–[121], resulting in increased DCS. However, dynamically increasing frequency in the ranges used here did not increase the firing of most excitatory neurons (Fig. 2,3) and if anything, likely drives stronger inhibition during the offset period due to preferential activation of PV neurons. However, at lower frequencies, dynamic modulation could result in better modulation of excitatory neurons and also less recruitment of PV-type neurons, although lower frequencies have other undesirable effects such as decreased recruitment and electrode-specific perceptual effects [10]. Altogether then, biomimetic frequency modulation is unlikely to reduce inhibitory recruitment and increase excitatory neuron activity during offset in parameter ranges that will be useful for clinical applications. However, lower frequencies can reduce the intensity of activation of neurons during the extended hold period. While frequency modulation of ICMS then can reduce DCS and impact individual neurons responses, dynamic frequency modulation is less powerful than dynamic amplitude modulation for evoking onset/offset responses. Further studies with larger frequency ranges can help us more fully understand the potential utility of frequency in ICMS applications and further our understanding of how neurons respond to ICMS.

### Limitations

We aimed in this study to evaluate biomimetic stimulus encoding paradigms within parameter ranges used in previous work in animals and humans [7], [8], [10], [34]–[36], [67]–[69], [79], which we expect to inform parameter selection and train design in future studies. Our approach allowed us to study the spatiotemporal activation and DCS/SIDNE of cortical neurons, but provided many important limitations. First, we investigated these stimulus trains in mice. While the approach used in this paper is currently impossible in humans and highly difficult in primates, because rodent cortex is different than primate cortex, there may be overall differences in the response. However, while there are some important differences between species, cellular composition and density and cortical organization is similar across rodent and primate sensory cortices [122]–[125]. Therefore, this work is still highly informative to the basic mechanisms of ICMS, as rodent work has been useful in the past for understanding ICMS [34], [35], [37], [38], [77], [126] as well as the basic science of sensory cortices [30], [103], [108], [110], [127]. Effects in humans will still need to be evaluated, although high through-put work done in rodent models should also allow for lower sample sizes in humans based on a deeper understanding of the basic science. We believe that combining these approaches with cross-species analysis will yield the best progress in ICMS applications.

Another limitation is that we targeted a single plane within layer 2/3. Imaging deeper in the brain with high-resolution is currently difficult due to light scattering and sparse GCaMP expression in layer 4 [128]. However, with new emerging technologies, we hope to expand our work to this layer as well. Currently, ICMS is targeted to layer 4 for sensory restoration, although in humans electrode location is impossible to verify following implant, so that some electrode implants may be in layer 2/3 or the array may be slanted and electrodes on a single array may span multiple layers. Differences in layers activated could result in electrode specific effects of ICMS [129], [130]. Therefore, while stimulation might be ideally targeted to layer 4, this picture is still not clear, and stimulation in all layers will be informative to future approaches for ICMS.

The temporal resolution of GCaMP6s also limits our ability to measure certain properties of neural activation. Previous studies have reported that GCaMP6s has latencies of 300-500 ms [82], [83]. Indeed, we saw here that the peak of activation for short trains occurred around 500 ms, where we would expect the peak to be around 0- 200 ms, aligning with this previous work. Because of these limitations in GCaMP, we are unable to evaluate the entrainment of the neurons to different frequencies. While explicit firing rates are difficult to extract, increases or decreases in calcium evoked activity do represent real changes in firing rate (i.e. no calcium activity means no action potentials). Given these limitations, we focused our investigation on the spatiotemporal relationships and DCS/SIDNE of neurons over time. Latencies in GCaMP may have influenced Short train responses, where detected facilitation for Short trains is actually due to overlapping calcium evoked by each transient. In this scenario, we were careful to recognize latencies as a possible influence. Therefore, these limitations do not impact our reported findings. For mouse experiments, genetically encoded voltage indicators (GEVI) are an alternative approach to Ca2+ sensors. GEVIs would be preferred due to direct visualization of ICMS evoked potentials; however, their development has lagged relative to Ca2+ sensors and suffer from low SNR making GEVIs difficult for *in vivo* use. Another difficulty lies with the extra-cellular positioning of the current GEVI sensors, which make delineation of individual cells from nearby neurites difficult [131]. If these technologies are improved in the future, then GEVIs could be used to look at the same dynamics with better temporal resolution.

These studies were also conducted in anesthetized animals. Cortical activity during wakefulness may impact the response of ICMS [79]. While we expect the influence on the local recruited population to be minimal, wakefulness can impact downstream activation. Future studies should evaluate any differences between the activation with ICMS in anesthetized vs. awake preparations to understand how these effects may differ.

Finally, we were only able to test a limited set of parameters because of the time limitations of the study (3-5 hour) and the theoretically infinite ICMS parameter space and many possible combinations of the selected parameters. The parameters used here were sufficient for the analyses and conclusions we provide. Short trains were designed to inject the same amount of charge, which allowed us to control effects driven by total charge injection and ensure that the effects observed were driven by differences in the distribution of charge throughout the trains. Frequency was matched between BioAmp and Fixed trains and frequency was matched between BioBoth and BioFreq trains, so differences in the evoked activity between these pairs could only be related to dynamic changes in the amplitude. While neurons may have been better able to entrain to lower frequency ranges, here we focused on high frequency ranges since these ranges are reflective of relevant ranges used in human studies. Future studies then should test other combinations of ICMS parameters, particularly other frequency ranges, to better understand how to optimally encode stimulus input.

### Conclusion: Implications for brain-computer interfaces

ICMS is an emerging approach to restore sensations, primarily somatosensory and visual. Traditionally, researchers have encoded sensory input directly with linear/engineered approaches. Here, biomimetic amplitude modulation evoked strong onset and offset responses. Improvements in robotic arm control with sensory feedback happen primarily around initial object contact, implying that initial confidence in sensory input is critical for improved robotic arm control [41]. We also found that all trains resulted in significant success in intensity discrimination for an ideal observer, which agree with object identification and amplitude discrimination results in humans [60], [62]. Here, we applied these trains in both visual and somatosensory cortices. Although onset and offset activity may be more traditionally associated with somatosensation, onset/offset has also been observed in the visual system [45], [47]–[50]. However, there are clearly differences between somatosensory and visual cortices and the demands of each of these modalities. Activation of neurons was similar across these two cortices, but if these paradigms will be equally effective for perception in both modalities is unclear.

Here, we showed biomimetic amplitude modulation maintained excitability over long periods of stimulation and reduced the total charge injection. Biomimetic frequency modulation in these clinically relevant parameter ranges only modulated a small number of neurons, but was also able to reduce depression to continuous ICMS relative to Fixed trains that had higher frequencies overall. BioBoth trains were dominated by amplitude effects so that the effects were similar to BioAmp trains. One notable difference between BioAmp and BioBoth trains was the intensity and depression induced by Short trains, where BioBoth evoked lower intensities and more depression. This could be because the short break between the onset/offset had a larger impact on the effects, and high-amplitude and high-frequency transients depressed the network more. The interaction then between amplitude, frequency, and duration in evoked responses and depression should be studied in more detail in future work. However, decreases in frequency may also allow for decreased stimulation artifact, which can still be useful for clinical applications [61]. Because continuous stimulation, especially at high amplitudes or frequencies, can lead to electrode degradation [132], [133], reducing charge injection may improve the longevity of implanted arrays. While parameters can be arbitrarily selected to result in more or less charge injection, any reasonable set of ICMS parameters should result in less charge injection for biomimetic encoding schemes vs. fixed encoding schemes. Indeed, decreases in charge injected has been previously noted as a potential strength for this method of encoding [60]. This simple modification to ICMS encoding then may provide many benefits for clinical applications.

## Supporting information

Supplemental Movie 1: Short BioAmp train activation in Mouse 2. Recorded calcium responses of neurons to BioAmp Short trains in Mouse 2.

Supplemental Movie 2: Short BioBoth train activation in Mouse 2.

Supplemental Movie 3: Short BioFreq train activation in Mouse 2.

Supplemental Movie 4: Short Fixed train activation in Mouse 2.

Supplemental Movie 5: Long BioAmp train activation in Mouse 7.

Supplemental Movie 6: Long BioBoth train activation in Mouse 7.

Supplemental Movie 7: Long BioFreq train activation in Mouse 7.

Supplemental Movie 8: Long Fixed train activation in Mouse 7.

## Acknowledgements

The authors would like to thank Kevin Stieger, Keying Chen, Naofumi Suematsu, Fan Li, and Adam Forrest for feedback. This work was supported by NIH T32 NS086749, R01NS105691, R01NS115707 and NSF CAREER CBET 1943906.

---

**Supplementary Table 1:** **Mice used for all experiments and corresponding trains presented/area stimulated.** Mice are named based on the chronological order in which the animals were tested. Short trains represent the 1-s on 4-s off paradigm. Long trains represent the 30-s on 15-s off paradigm. Long trains+ represent the 30-s on 15-s off paradigm at an elevated amplitude.

|  | Short trains | Long trains | Long trains+ | Cortical area | Sex |
| --- | --- | --- | --- | --- | --- |
| **Mouse 1** | 2 | 0 | 0 | Visual cortex | Male |
| **Mouse 2** | 2 | 0 | 0 | Visual cortex | Female |
| **Mouse 3** | 2 | 0 | 0 | Visual cortex | Female |
| **Mouse 4** | 2 | 2 | 0 | Visual cortex | Male |
| **Mouse 5** | 2 | 2 | 0 | Visual cortex | Female |
| **Mouse 6** | 1 | 1 | 1 | Visual cortex | Female |
| **Mouse 7** | 1 | 1 | 1 | Somatosensory cortex | Male |
| **Mouse 8** | 1 | 1 | 1 | Somatosensory cortex | Male |
| **Mouse 9** | 1 | 1 | 1 | Somatosensory cortex | Female |
| **Mouse 10** | 1 | 1 | 1 | Visual cortex | Male |
| **Mouse 11** | 1 | 1 | 1 | Somatosensory cortex | Female |
| **Mouse 12** | 1 | 1 | 1 | Visual cortex | Male |

**SuppMov1_BioAmp_Short.avi**

**Supplemental Movie 1: Short BioAmp train activation in Mouse 2.** Recorded calcium responses of neurons to BioAmp Short trains in Mouse 2. Time series were processed with SVD to improve clarity. 30 Hz images were group averaged into 10-frame averages resulting in 3 Hz. Videos were exported at 6 fps, representing a 2x playback. “Stim ON” in the upper left corner indicates when ICMS was being delivered. The total field size is 543.2×543.2 µm

**SuppMov2_BioBoth_Short.avi**

**Supplemental Movie 2: Short BioBoth train activation in Mouse 2.** Recorded calcium responses of neurons to BioBoth Short trains in Mouse 2. Time series were processed with SVD to improve clarity. 30 Hz images were group averaged into 10-frame averages resulting in 3 Hz. Videos were exported at 6 fps, representing a 2x playback. “Stim ON” in the upper left corner indicates when ICMS was being delivered. The total field size is 543.2×543.2 µm

**SuppMov3_BioFreq_Short.avi**

**Supplemental Movie 3: Short BioFreq train activation in Mouse 2.** Recorded calcium responses of neurons to BioFreq Short trains in Mouse 2. Time series were processed with SVD to improve clarity. 30 Hz images were group averaged into 10-frame averages resulting in 3 Hz. Videos were exported at 6 fps, representing a 2x playback. “Stim ON” in the upper left corner indicates when ICMS was being delivered. The total field size is 543.2×543.2 µm

**SuppMov4_Fixed_Short.avi**

**Supplemental Movie 4: Short Fixed train activation in Mouse 2.** Recorded calcium responses of neurons to Fixed Short trains in Mouse 2. Time series were processed with SVD to improve clarity. 30 Hz images were group averaged into 10-frame averages resulting in 3 Hz. Videos were exported at 6 fps, representing a 2x playback. “Stim ON” in the upper left corner indicates when ICMS was being delivered. The total field size is 543.2×543.2 µm

**SuppMov5_BioAmp_Long.avi**

**Supplemental Movie 5: Long BioAmp train activation in Mouse 7.** Recorded calcium responses of neurons to BioAmp Long trains in Mouse 7. Time series were processed with SVD to improve clarity. 30 Hz images were group averaged into 8-frame averages resulting in 3.25 Hz. Videos were exported at 7.5 fps, representing a 2x playback. “Stim ON” in the upper left corner indicates when ICMS was being delivered. The total field size is 543.2×543.2 µm

**SuppMov6_BioBoth_Long.avi**

**Supplemental Movie 6: Long BioBoth train activation in Mouse 7.** Recorded calcium responses of neurons to BioBoth Long trains in Mouse 7. Time series were processed with SVD to improve clarity. 30 Hz images were group averaged into 8-frame averages resulting in 3.25 Hz. Videos were exported at 7.5 fps, representing a 2x playback. “Stim ON” in the upper left corner indicates when ICMS was being delivered. The total field size is 543.2×543.2 µm

**SuppMov7_BioFreq_Long.avi**

**Supplemental Movie 7: Long BioFreq train activation in Mouse 7.** Recorded calcium responses of neurons to BioFreq Long trains in Mouse 7. Time series were processed with SVD to improve clarity. 30 Hz images were group averaged into 8-frame averages resulting in 3.25 Hz. Videos were exported at 7.5 fps, representing a 2x playback. “Stim ON” in the upper left corner indicates when ICMS was being delivered. The total field size is 543.2×543.2 µm

**SuppMov8_Fixed_Long.avi**

**Supplemental Movie 8: Long Fixed train activation in Mouse 7.** Recorded calcium responses of neurons to Fixed Long trains in Mouse 7. Time series were processed with SVD to improve clarity. 30 Hz images were group averaged into 8-frame averages resulting in 3.25 Hz. Videos were exported at 7.5 fps, representing a 2x playback. “Stim ON” in the upper left corner indicates when ICMS was being delivered. The total field size is 543.2×543.2 µm

**Supplementary Figure 1:**
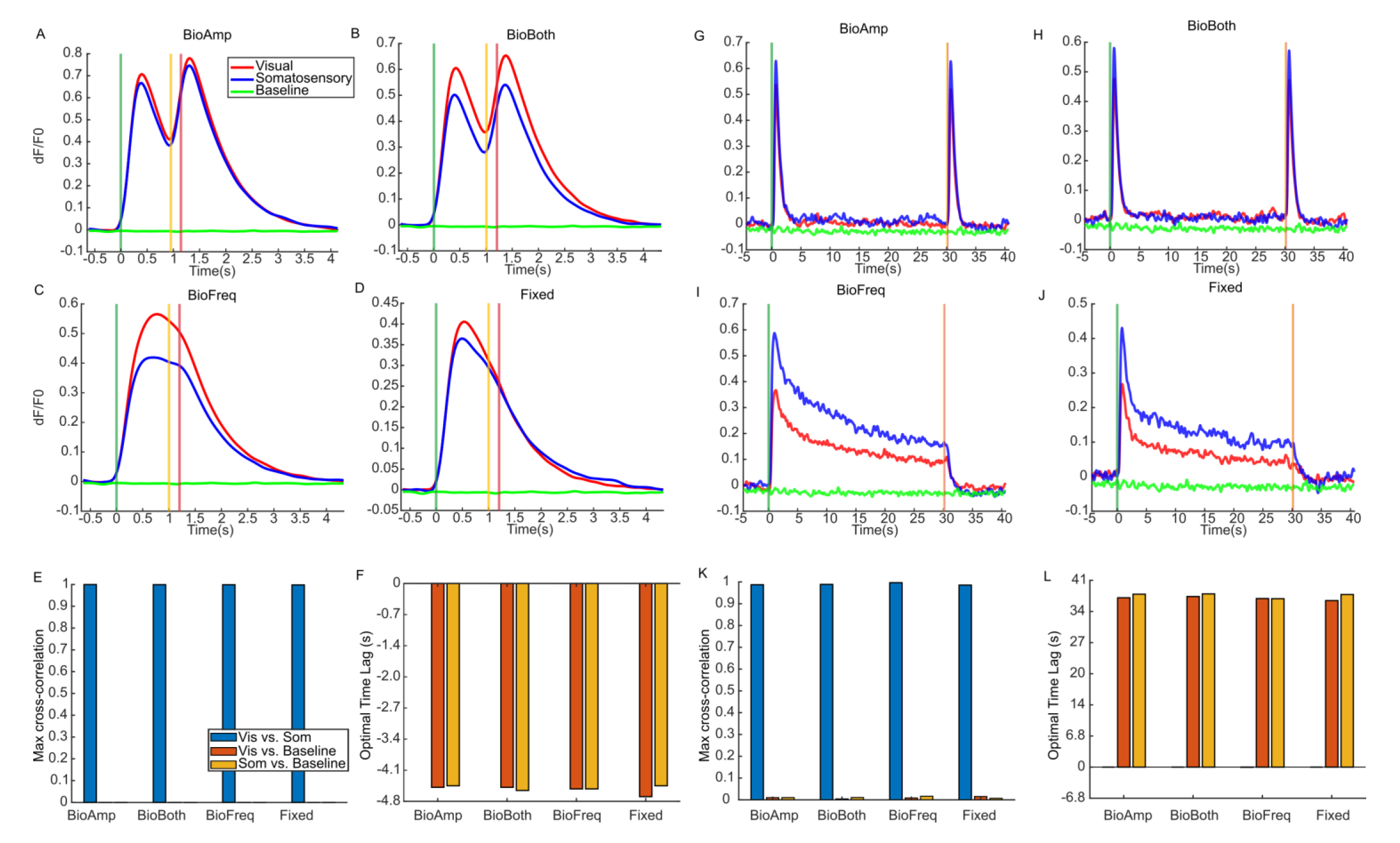
Responses are similar across somatosensory and visual cortices. A-D) Average traces for Short trains plotted for visual cortex (*n*=8), somatosensory cortex (*n*=4), and baseline (no ICMS, *n*=10). E) The max correlation value for cross-correlation between somatosensory and visual cortex, visual cortex and baseline, and somatosensory cortex and baseline for Short trains. The correlation value of near 1 shows a strong correlation between the evoked responses between visual and somatosensory cortex. F) The optimal lag for cross-correlations for Short trains (the lag between time series that maximizes the correlation). The value near 0 shows a strong temporal overlap in the evoked responses between visual and somatosensory cortex. G-J) Average traces for Long trains plotted for visual cortex (*n*=5), somatosensory cortex (*n*=4), and baseline (no ICMS, *n*=7). K) The max correlation value for cross-correlation for Long trains. L) The optimal lag for cross-correlations for Long trains.

**Supplementary Figure 2:**
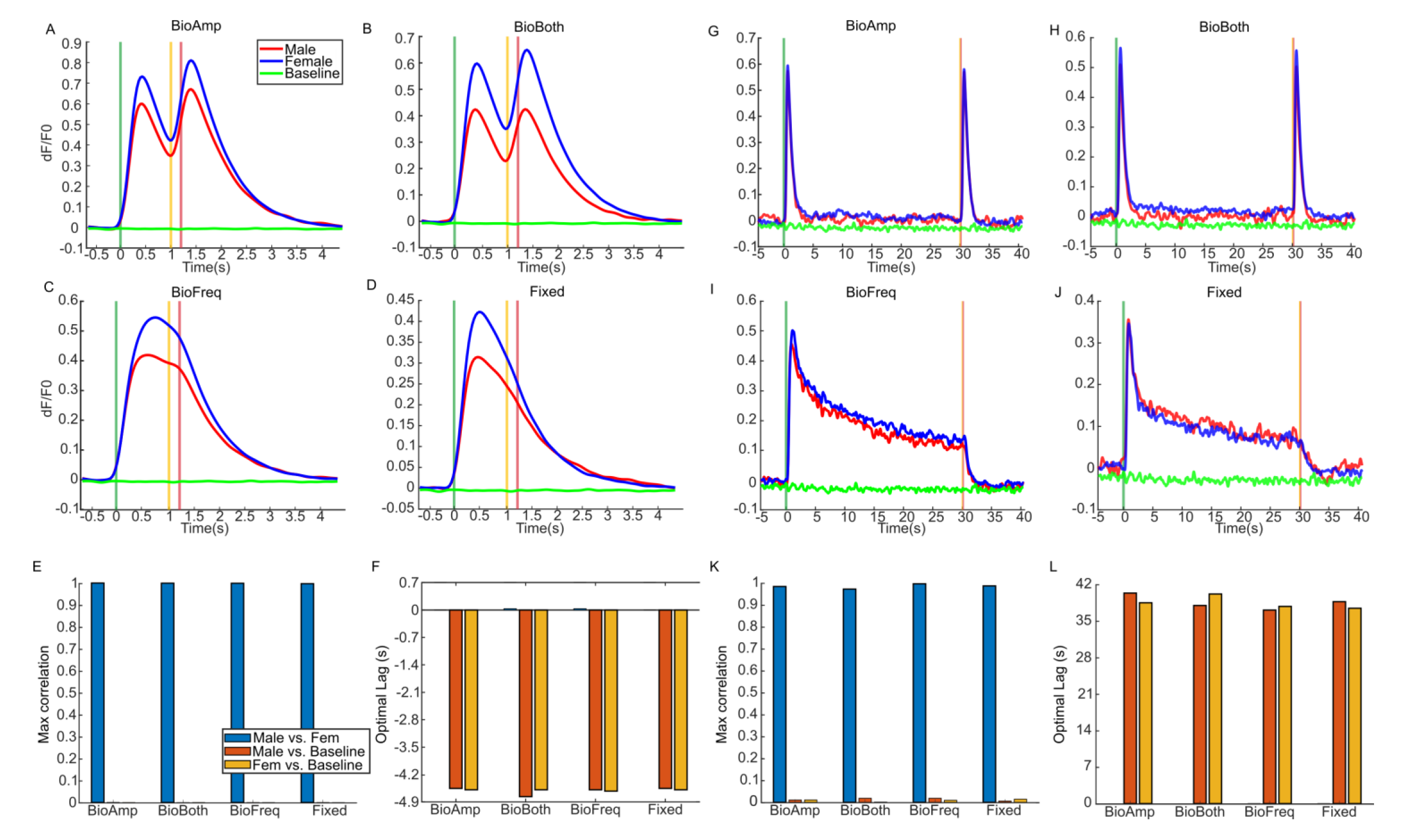
Responses are similar across animal sex. A-D) Average traces for Short trains plotted for males (*n*=6), females (*n*=6), and baseline (no ICMS, *n*=10). E) The max correlation value for cross-correlation between males and females, males and baseline, and females and baseline for Short trains. The correlation value of near 1 shows a strong correlation between the evoked responses between visual and somatosensory cortex. F) The optimal lag for cross-correlations for Short trains (the lag between time series that maximizes the correlation). The value near 0 shows a strong temporal overlap in the evoked responses between visual and somatosensory cortex G-J) Average traces for Long trains plotted for males (*n*=5), females (*n*=4), and baseline (no ICMS, *n*=7). K) The max correlation value for cross-correlation for Long trains. L) The optimal lag for cross-correlations for Long trains.

**Supplementary Figure 3:**
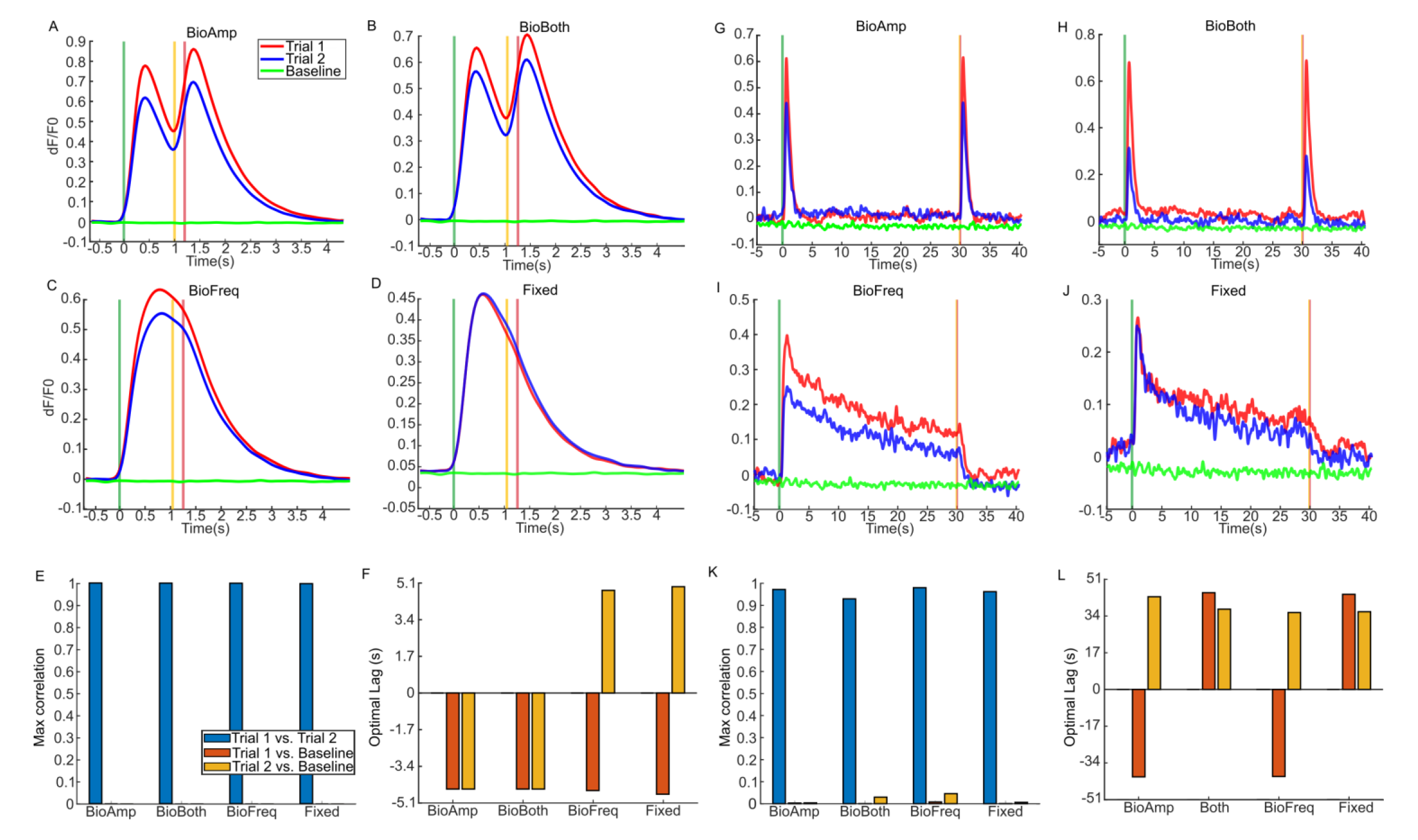
Responses are similar across trials. A-D) Average traces for Short trains plotted for trial 1 (*n*=5), trial 2 (*n*=5), and baseline (no ICMS, *n*=10). E) The max correlation value for cross-correlation between trial 1 and trial 2, trial 1 and baseline, and trial 2 and baseline for Short trains. The correlation value of near 1 shows a strong correlation between the evoked responses between visual and somatosensory cortex. F) The optimal lag for cross-correlations for Short trains (the lag between time series that maximizes the correlation). The value near 0 shows a strong temporal overlap in the evoked responses between visual and somatosensory cortex G-J) Average traces for Long trains plotted for trial 1 (*n*=2), trial 2 (*n*=2), and baseline (no ICMS, *n*=7). K) The max correlation value for cross-correlation for Long trains. L) The optimal lag for cross-correlations for Long trains.

**Supplementary Figure 4:**
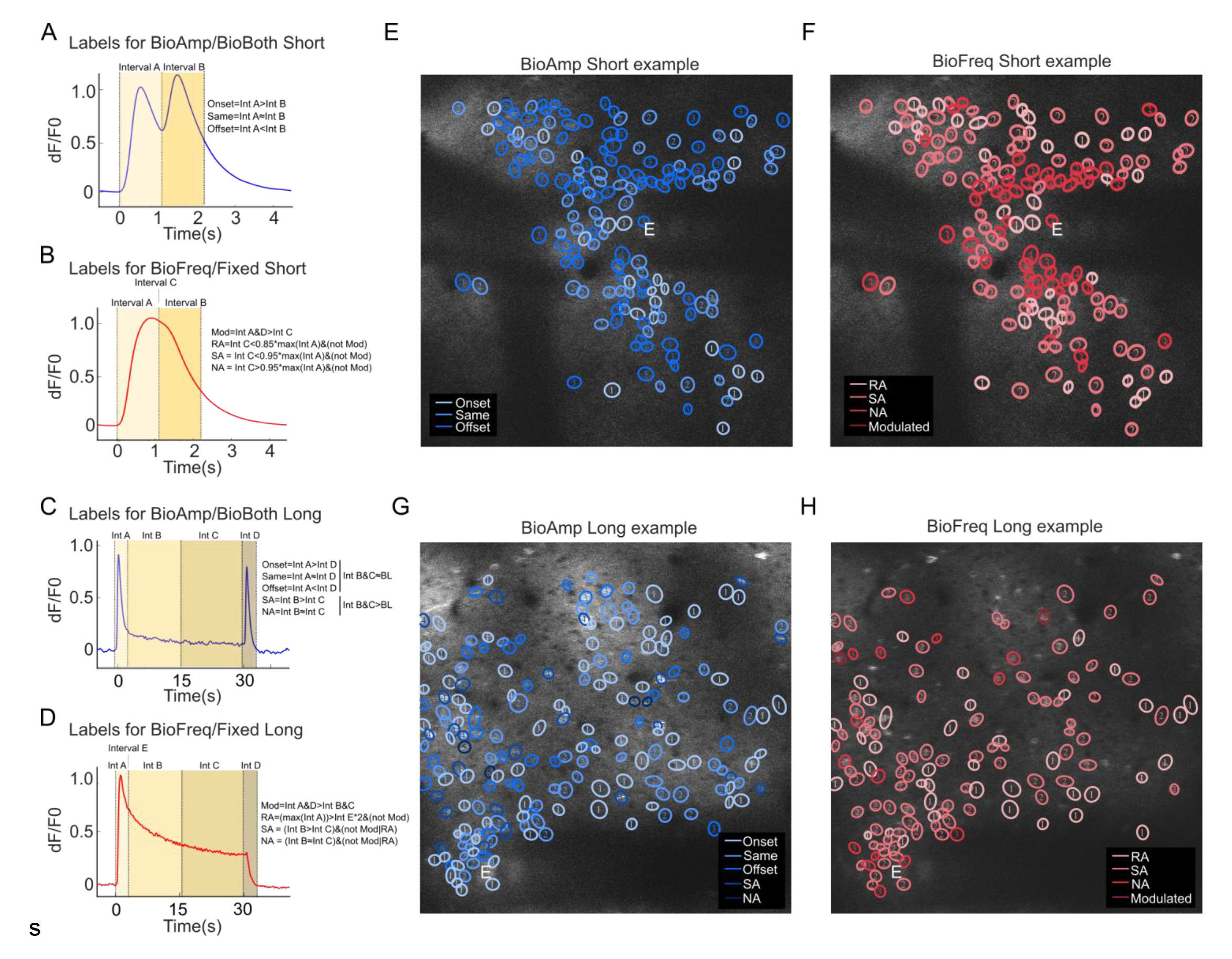
Neurons were divided based on identified temporal characteristics. A) Short BioAmp and BioBoth trains were identified to have differences in the location of maximum peak. If the first peak was greater than the second peak by at least 0.5x standard deviations of the noise, then the neuron was identified as “Onset.” If the second peak was greater, the neuron was identified as “Offset.” If the peaks were not different by more than 0.5x standard deviations of the noise, then the neuron was identified as “Same.” B) Short BioFreq trains were identified to have a small number of neurons that were modulated by the onset and offset transients. These “Modulated” neurons were separated if they had maximum intensities during the Onset and Offset period that was greater than the Hold period by more than 0.5x standard deviations of the noise. Neurons that did not have this difference were classified “Unmodulated.” All Fixed trains were identified as “Unmodulated.” C) Long and Long+ BioAmp and BioBoth trains were identified similarly with “Onset,” “Same,” and “Offset,” but additionally had some neurons that had elevated activity during the hold period, which were separated as “Slowly Adapting” (SA) or “Non-adapting” (NA). SA neurons had a maximum intensity in the second half of the hold period that was less than the first half by at least 0.5x standard deviations while NA neurons had a maximum in the second half that was not different than the first half by at least 0.5x standard deviations. D) Long and Long+ BioFreq and Fixed trains similarly had SA and NA responses, but also had “Rapidly Adapting” (RA) responses. These RA responses were characterized by the maximum intensity at 10-s after stimulation decreasing to less than half of the maximum intensity within the first 10-s. E-H) Examples of active neurons and their classes for a BioAmp Short (E), BioFreq Short (F), BioAmp Long+ (G) and BioFreq Long+ (H) trains. The white letter E indicates the electrode location. Each circle shows a manually labeled neuron that was exported from ImageJ and fit with an ellipse in MATLAB [134]. Colors and numbers indicate the class of each neuron.

**Supplementary Figure 5:**
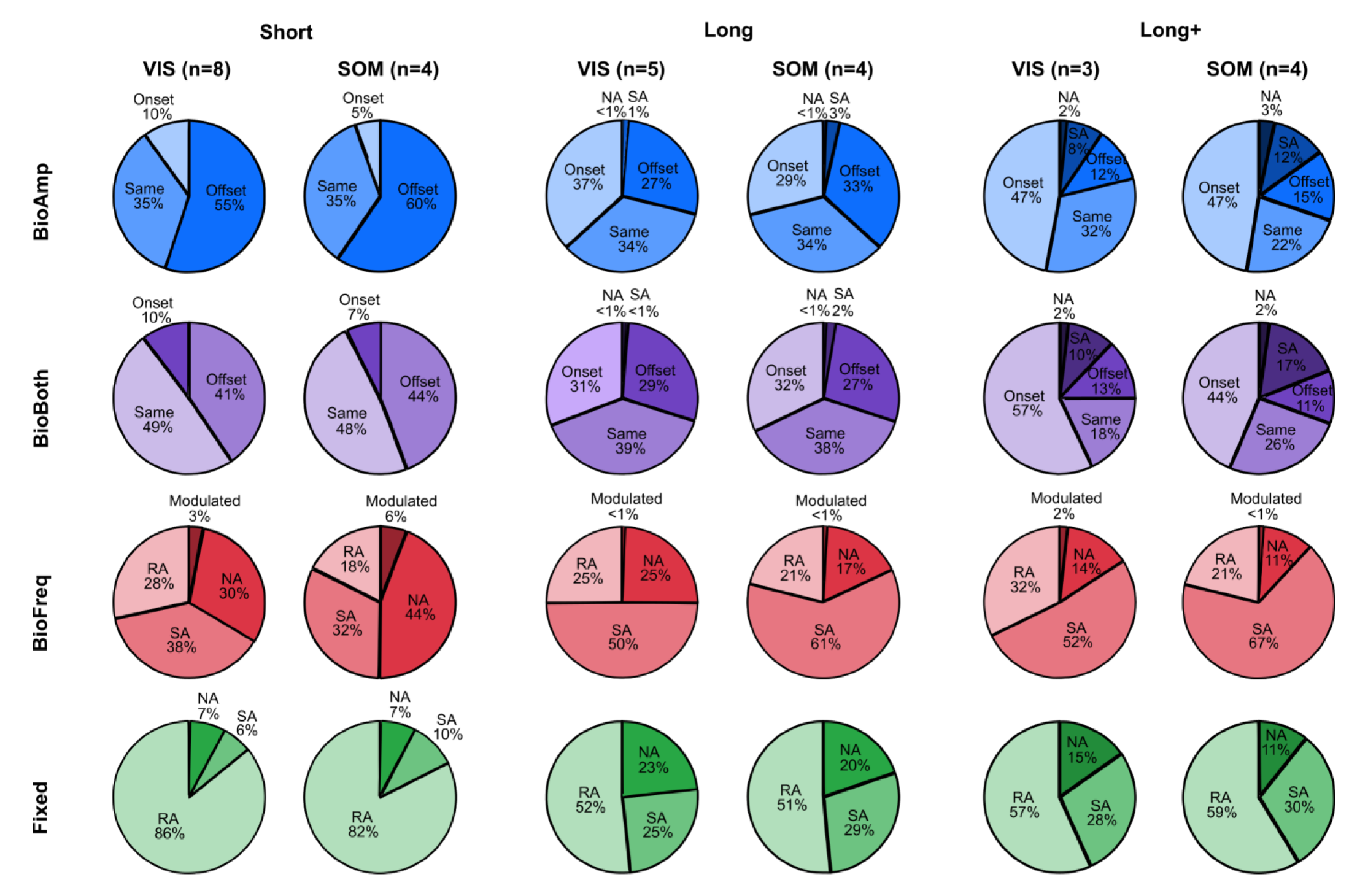
Distribution of classes is similar across visual and somatosensory cortices. Pie plots showing the percentage of each class for each train type (BioAmp, BioBoth, BioFreq, and Fixed) as well as each train length (Short, Long, Long+). For each train type, the distributions of each class are similar for visual (VIS) and somatosensory (SOM) cortices.

**Supplementary Figure 6:**
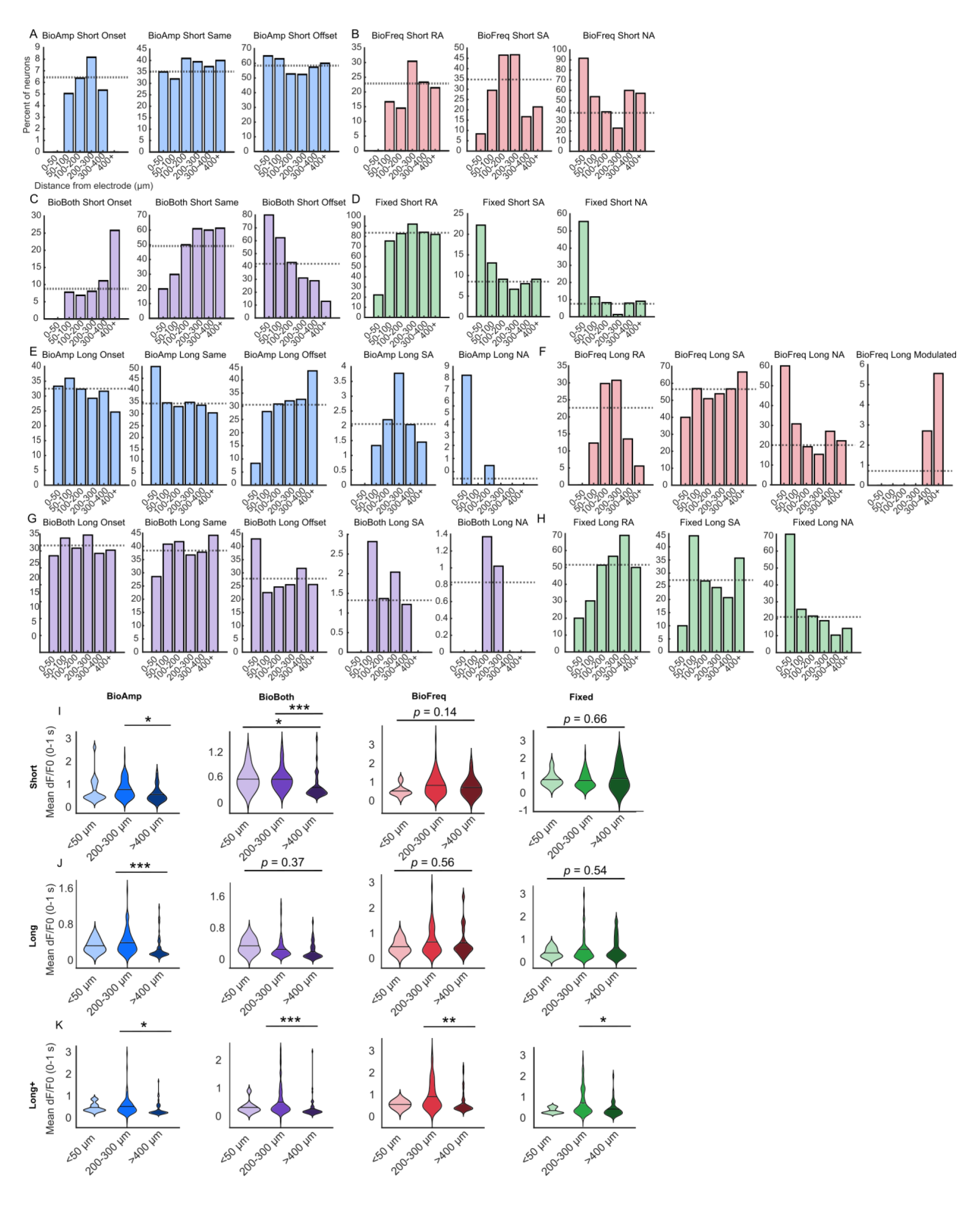
Relationship between distance from the electrode and neuronal class across all train types. Each bar shows the percent of neurons at the given distance from the electrode that were identified as the label given in the title. The horizontal line shows the average percent of neurons across all distances with the given label. Percents are within distance bins and not within label, so bars within a plot will not add up to 100%, but bars across plots will. For example, in A, adding up the 0-50 bar in all three plots will add to 100%. A-D) Short train responses for BioAmp (A), BioFreq (B), BioBoth (C), and Fixed (D) trains. Generally, neurons closer to the electrode were more likely to be Offset (A,C) or Non-adapting (B,D) indicating stability or facilitation, while neurons farther were more likely to be Onset (A,C) or Rapidly adapting (B,D) indicating depression. E-H) Long train responses for BioAmp (E), BioFreq (F), BioBoth (G), and Fixed (H) trains. Neurons closer to the electrode were more likely to be NA and neurons farther were more likely to be RA, but the relationships were weak overall for the BioAmp and BioBoth trains. I-K) Violin plots comparing the mean intensity across distances within the first 1 s of ICMS for Short (I), Long (J), and Long+ (K) trains. For BioAmp and BioBoth trains, neurons closer to the electrode tended to have higher intensities (significant for Short trains). For BioFreq and Fixed trains, neurons farther from the electrode tended to have higher intensities (significant for Long and Long+ trains).

**Supplementary Figure 7:**
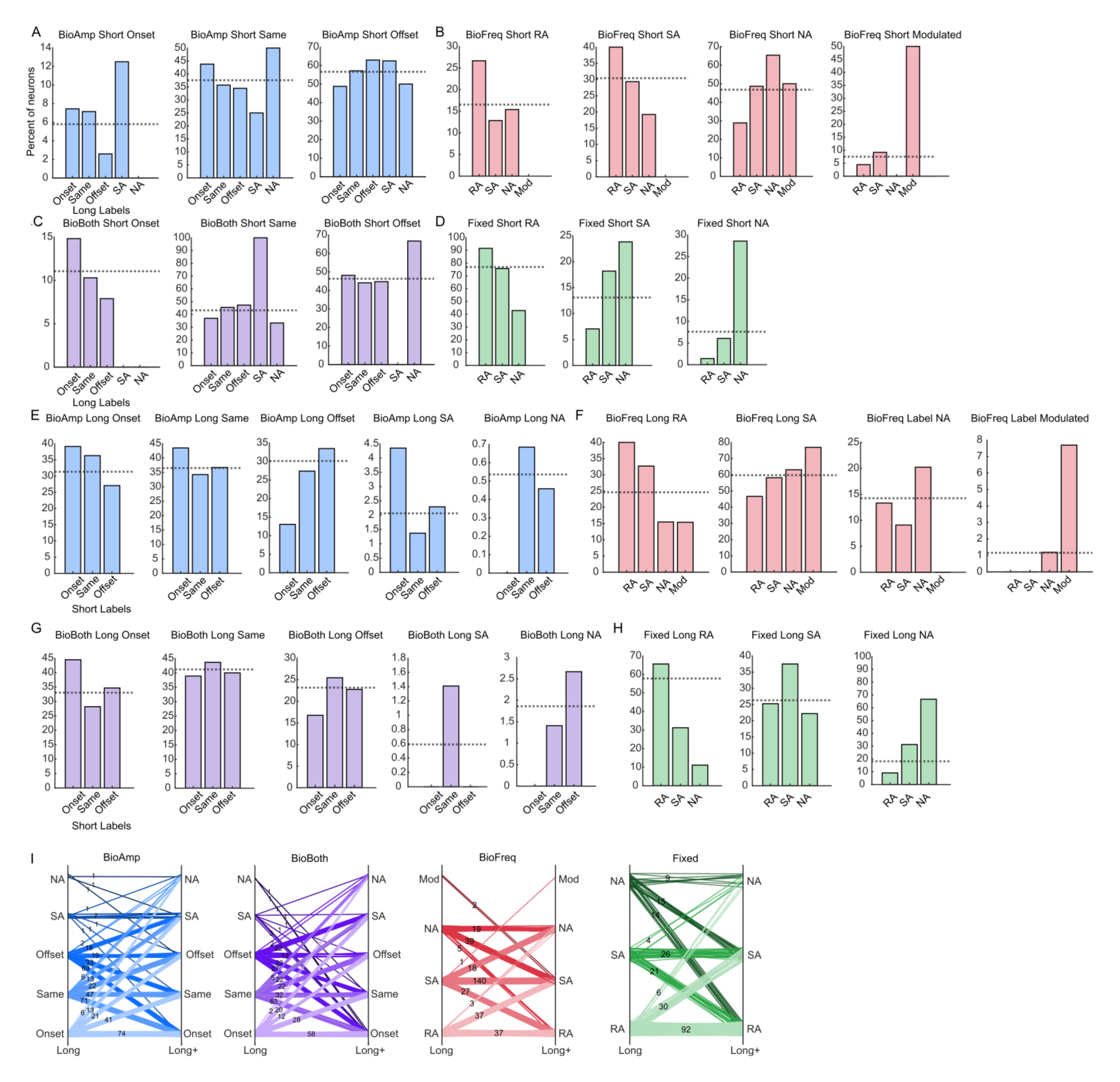
Relationship between classes evoked by Short and Long trains. Each bar shows the percent of neurons with the label given on the x-axis that were identified as the label given in the title. The horizontal line shows the average percent of neurons across all neurons with the label given in the title. Percentsw are within the labels in the x-axis, so bars within a plot will not add up to 100%, but bars across plots will. For example, in A, adding up the Onset bar in all three plots will add to 100%. A-D) Long train classes grouped by Short train classes. for BioAmp (A), BioFreq (B), BioBoth (C), and Fixed (D) trains. Generally, neurons identified with one class with the Short trains were more likely to be identified with the same class for the Long train. This relationship was weaker for the BioAmp and BioBoth trains, which is likely related to the increase in classes for Long trains (SA and NA) as well as the reduction in adaptation relative to BioFreq and Fixed train (because the class is strongly related to rate and magnitude of depression) E-H) Short train classes grouped by Long train classes for BioAmp (E), BioFreq (F), BioBoth (G), and Fixed (H) trains. Neurons identified with one class with the Short trains were more likely to be identified with the same class for the Long train. This relationship was again weaker for BioAmp and BioBoth trains. I) Parallel plots showing how neurons change classes between Long and Long+ trains. Generally, neurons were more likely to be depressed (Onset or RA) for the Long+ train and more likely to show increased activity during the hold phase for BioAmp and BioBoth trains.

**Supplementary Figure 8:**
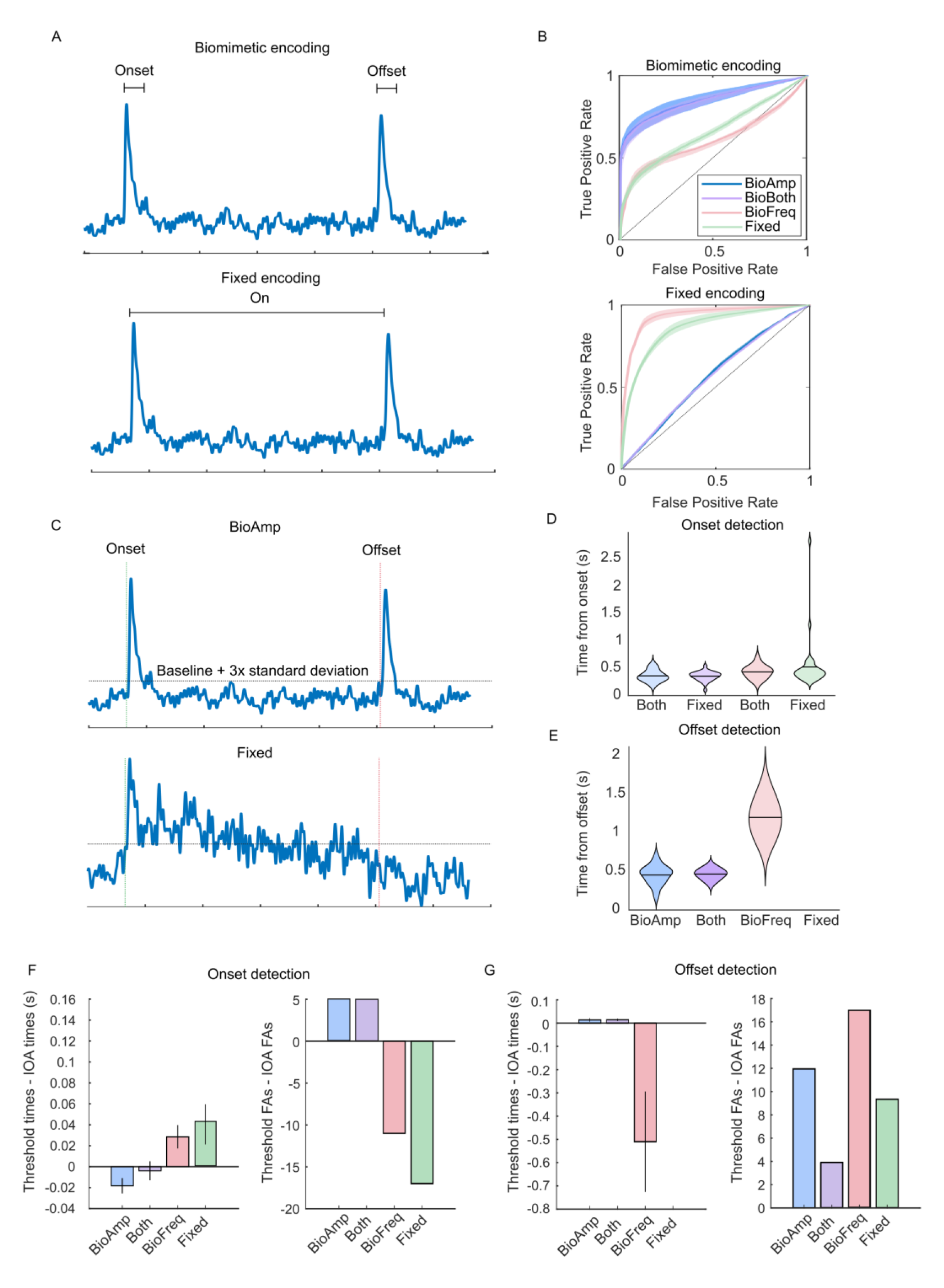
Encoding training schemes for ideal observer analysis. A) We used two methods of encoding: biomimetic (top) or fixed (bottom). For biomimetic encoding, onset and offset periods were labeled and input to the classifier. For fixed encoding the total on period (onset and the hold period) were labeled and input to the classifier. Here, those periods are marked on an example BioAmp trace. The inputs were dF/F_0_ profiles and corresponding labels of the stimulus state. Onset and offset detection times (DTs) were the times classifier scores reached 75% confidence in the predicted label. A false positive was when 75% confidence was achieved before the known onset or offset, and trials with false positives were excluded from calculation of average onset and offset times. B) Performance of each classifier for each train type assessed with a Receiver Operator characteristic. <u>The</u> BioAmp and BioBoth trains had a larger area under the curve for the biomimetic encoding scheme, while the Fixed and BioFreq trains had a larger area under the curve for the fixed encoding scheme. C) Using just the Baseline + 3x standard deviation to determine the onset/offset or on period for an example BioAmp (top) and example Fixed (bottom) train. D) Onset detection for each train using the simple baseline+noise threshold. E) Offset detection for each train using the simple baseline+noise threshold. Note that there is no mean for Fixed because all Fixed trains resulted in early detection of offset due to the adapting responses (false alarm). F) Comparison of onset determination with the simple threshold method and ideal observer analysis. Onset detection was slightly faster for BioAmp and BioBoth trains and slower for BioFreq and Fixed trains with IOA. IOA resulted in less false alarms for BioAmp and BioBoth trains but more false alarms for BioFreq and Fixed trains. G) The simple threshold method resulted in lower offset times for BioFreq and similar times for BioAmp and BioFreq. The simple threshold method resulted in much more false alarms for offset across all trains.

